# Distinct systemic and gut IgA responses to bacteria of the human upper gastrointestinal tract

**DOI:** 10.1101/2025.07.01.662496

**Authors:** Filipa Vaz, Ida Lindeman, Henriette H. Hoffmann, Christine Skagen, Anne K. Steffensen, Jørgen V. Bjørnholt, Geir E. Tjønnfjord, Knut E. A. Lundin, Ludvig M. Sollid, Rasmus Iversen

**Affiliations:** Norwegian Coeliac Disease Research Centre, Institute of Clinical Medicine, University of Oslo, NO-0372 Oslo, Norway; Department of Immunology, Oslo University Hospital, NO-0372 Oslo, Norway; Department of Microbiology, Oslo University Hospital, NO-0372 Oslo, Norway; Department of Haematology, Oslo University Hospital, NO-0372 Oslo, Norway; Department of Gastroenterology, Oslo University Hospital, NO-0372 Oslo, Norway

**Keywords:** Gut, duodenum, serum, bone marrow, plasma cells, IgA, bacteria, microbiota, commensal, Neisseria, IgA1 proteases

## Abstract

The mucosa lining the gastrointestinal tract harbors the body’s largest population of plasma cells, most of which produce dimeric IgA destined for release into the lumen. In addition, there is systemic production of monomeric IgA circulating in the blood. Little is known about the connection between systemic and mucosal IgA. To address this relationship and to explore antibody responses against the microbiota, we isolated bacteria from duodenal biopsies and assessed antibody reactivity. Systemic IgA showed reactivity to bacteria of the upper gastrointestinal tract with a preference for binding *Neisseria* species, while duodenal IgA showed broader reactivity. We found limited clonal overlap between gut and bone marrow plasma cells of individual donors, yet a few shared clones specific to bacterial antigens were identified. Despite showing clonal overlap, gut and bone marrow plasma cells have distinct IgA subclass distributions, and they likely depend on B-cell activation at discrete anatomical sites.

## INTRODUCTION

The intestinal mucosa is a major site of continuous antibody secretion by plasma cells. Most mucosal plasma cells produce immunoglobulin A (IgA)-class antibodies, which are secreted in the lamina propria as dimers connected via the joining (J) chain. Dimeric IgA is subsequently transported across the epithelium through association with the polymeric Ig receptor (pIgR) followed by release into the gut lumen together with a fragment of pIgR known as the secretory component.^1^ The resulting complex is termed secretory IgA, and its function is to bind luminal antigens of which bacteria make up a large proportion. Secretory IgA can confer protection against pathogens, and it has a fundamental role in controlling microbiota composition and interactions between bacteria. It can regulate bacterial phenotypes, modulate the production of certain metabolites, determine localization of bacteria within gut niches, and manage bacterial numbers.^2–5^

In addition to secretory IgA, there is systemic IgA circulating in blood. Unlike IgA produced at mucosal sites, serum IgA is monomeric, and it is believed to be produced mainly by bone marrow (BM) plasma cells.^6^ Mucosal immune responses in mice have been shown to give rise to BM plasma cells,^7–9^ but the relationship between mucosal and systemic IgA remains elusive. While both commensal and colitogenic intestinal bacteria are coated with secretory IgA at steady state,^10–12^ serum antibodies were initially thought only to target systemic pathogens.^13,14^ More recently, however, it has been demonstrated that serum antibodies can also recognize commensals and that serum IgA can protect against sepsis in the event of a breach in the mucosal barrier.^15–17^

In humans, there are two subclasses of IgA: IgA1 and IgA2. Whereas serum IgA is predominantly IgA1, mucosal IgA shows varying distribution between the subclasses along the gastrointestinal (GI) tract with a decreasing IgA1:IgA2 ratio from proximal to distal sites.^18,19^ This observation might suggest that systemic IgA mainly arises through immune reactions in the upper GI tract. To investigate this possibility and to study the relationship between mucosal and systemic IgA, we isolated bacteria from duodenal biopsies and assessed IgA reactivity in matched serum samples. Our results indicate that serum IgA selectively targets bacteria present in the upper GI tract (comprising the mouth, pharynx, esophagus, stomach, and duodenum). Species belonging to the genus *Neisseria* appear to play a prominent role in systemic IgA responses against the microbiota, while mucosal IgA recognizes a broader range of bacteria. Further, single-cell RNA sequencing (scRNA-seq) analysis of BM and duodenal plasma cells suggests that the IgA-expressing populations in these two compartments largely originate from separate immune reactions, despite showing a degree of clonal overlap. Collectively, our study highlights a role of bacteria in the upper GI tract in generating systemic IgA through a mechanism that is distinct from the formation of secretory IgA in the gut.

## RESULTS

### Serum IgA shows reactivity to bacteria isolated from duodenal biopsies

To assess IgA reactivity to the microbiota of the upper GI tract, we isolated bacteria from duodenal biopsies of four individuals who underwent endoscopy to test for celiac disease-associated mucosal changes (Table S1). The bacteria that were grown from the biopsy specimens likely include species present at the mucosal surface of the duodenum as well as species present at proximal sites, including the mouth and pharynx, which were picked up through the endoscopy procedure. In total, we established 175 strains that were used to test for antibody reactivity. The isolates belonged to ten different bacterial genera and one yeast represented by two isolates of the species *Candida albicans* (Fig. 1A and Table S2). The identified species are in agreement with the expected flora and have previously been found to be commonly present in the upper GI tract.^20–27^

**Figure 1.**
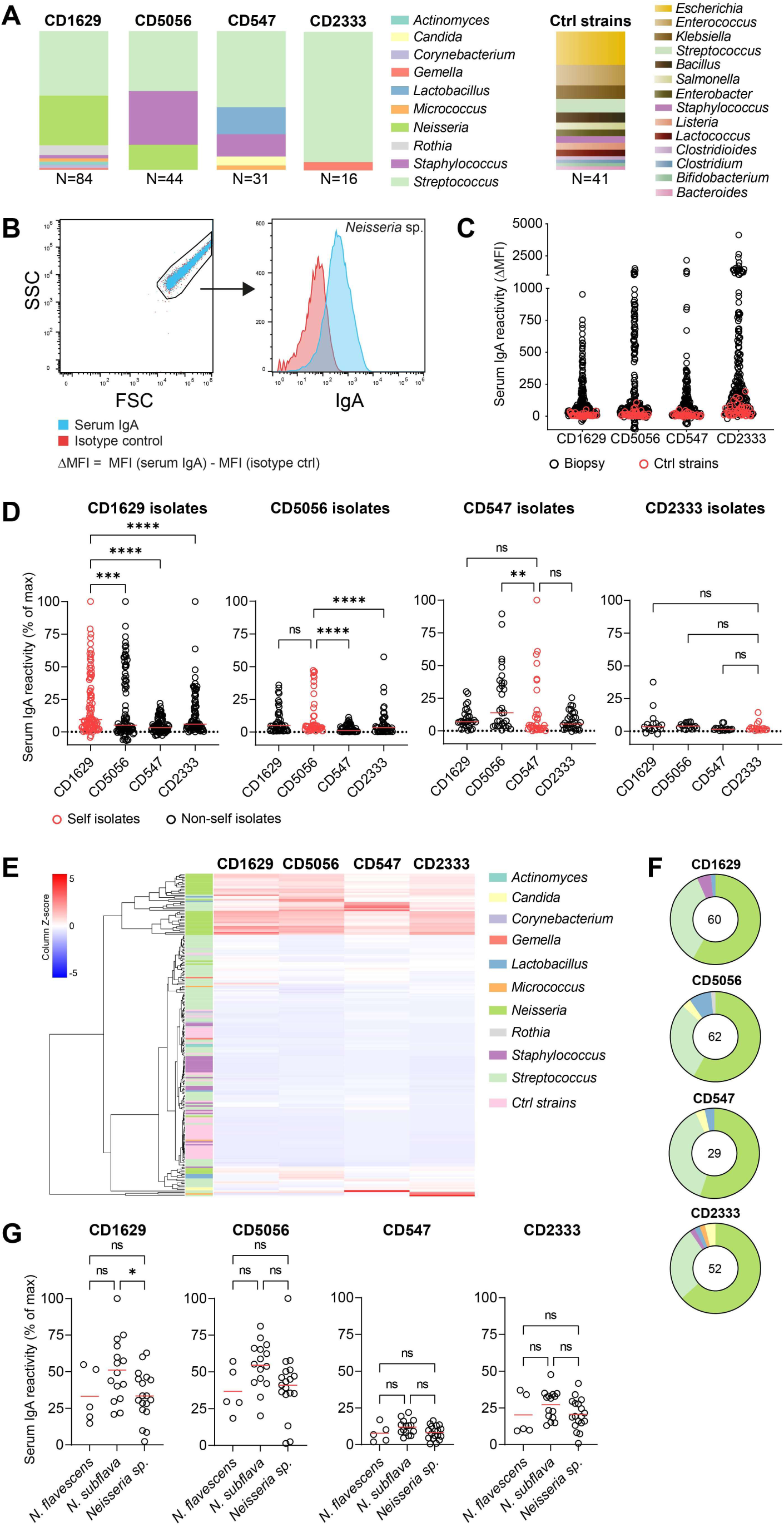
Serum IgA reactivity against bacteria. (A) Identified genera among all strains (N) isolated from duodenal biopsies of four donors as well as a panel of control reference strains. (B) Representative flow cytometry plots showing staining of a bacterial strain with purified serum IgA. (C) Reactivity of purified serum IgA of four donors against all biopsy isolates or control strains (n = 216) given as ΔMFI values determined by flow cytometry. (D) Serum IgA reactivity against strains isolated from each of the donors. Signals are given relative to the maximum ΔMFI value obtained for each IgA sample. Horizontal lines indicate medians, and difference between groups was evaluated by Friedman’s test with Dunn’s multiple comparisons correction. **p < 0.01, ***p < 0.001, ****p < 0.0001. (E) Heat map showing serum IgA reactivity of the four donors with all included strains. Each isolate (row) is labeled with its genus. (F) Distribution of IgA-reactive bacteria. For each serum sample (donor), only strains with ΔMFI values >10% of the maximum signal are displayed. Colors indicate genera as in (E). (G) Serum IgA reactivity against isolates of the genus *Neisseria*. The included isolates belonged to two different species or were unidentified (*Neisseria sp.*). Horizontal lines indicate means, and difference between groups was evaluated by one-way ANOVA with Holm-Sidak multiple comparisons correction. *p < 0.05.

We next purified serum IgA from blood samples of the same individuals and assessed binding to the whole, fixed bacteria by flow cytometry (Fig. 1B). We also included a panel of previously isolated control strains representing both pathogenic and non-pathogenic members of the microbiota, including species typically found in the colon. (Fig. 1A and Table S3). Serum IgA from each donor showed reactivity to several of the biopsy isolates, but we did not observe binding to the control strains (Fig. 1C). Some of the isolates were preferably recognized by donor-matching serum IgA, indicating a degree of privateness in systemic responses against the microbiota (Fig. 1D and Fig. S1). In general, however, the individual donors also showed reactivity to bacteria isolated from other donors. Thus, human serum IgA does not appear to specifically distinguish between “self” and “non-self” bacteria.

When analyzing serum IgA reactivity against the individual strains, we observed that strains belonging to the *Neisseria* genus were particularly well recognized across the four donors (Fig. 1E and 1F). Two different *Neisseria* species, *N. flavescens* and *N. subflava*, were identified in the panel. Both species were recognized by serum antibodies, but *N. subflava* isolates generally showed higher reactivity than *N. flavescens* isolates (Fig. 1G). Hence, serum antibodies might recognize both species-specific antigens and surface structures that are conserved within the *Neisseria* genus. Taken together, these results indicate that serum IgA selectively targets certain bacteria in the upper GI tract and that the antibodies effectively distinguish between different genera but not necessarily different species belonging to the same genus.

### Serum IgA reactivity to bacteria is stable over time

To test if systemic IgA responses against the microbiota vary over time, we took advantage of four serum samples collected from a single patient across a period of three years.

The purified IgA antibodies from each time point showed highly similar reactivity when assessing binding to our panel of bacteria (Fig. 2A and 2B). Among the IgA-reactive strains, *Neisseria* species gave particularly stable binding, whereas streptococci generally showed higher variability (Fig. 2C and Fig. S2). These results indicate that the systemic IgA response to bacteria of the upper GI tract is stable over time, particularly to *Neisseria* species.

**Figure 2.**
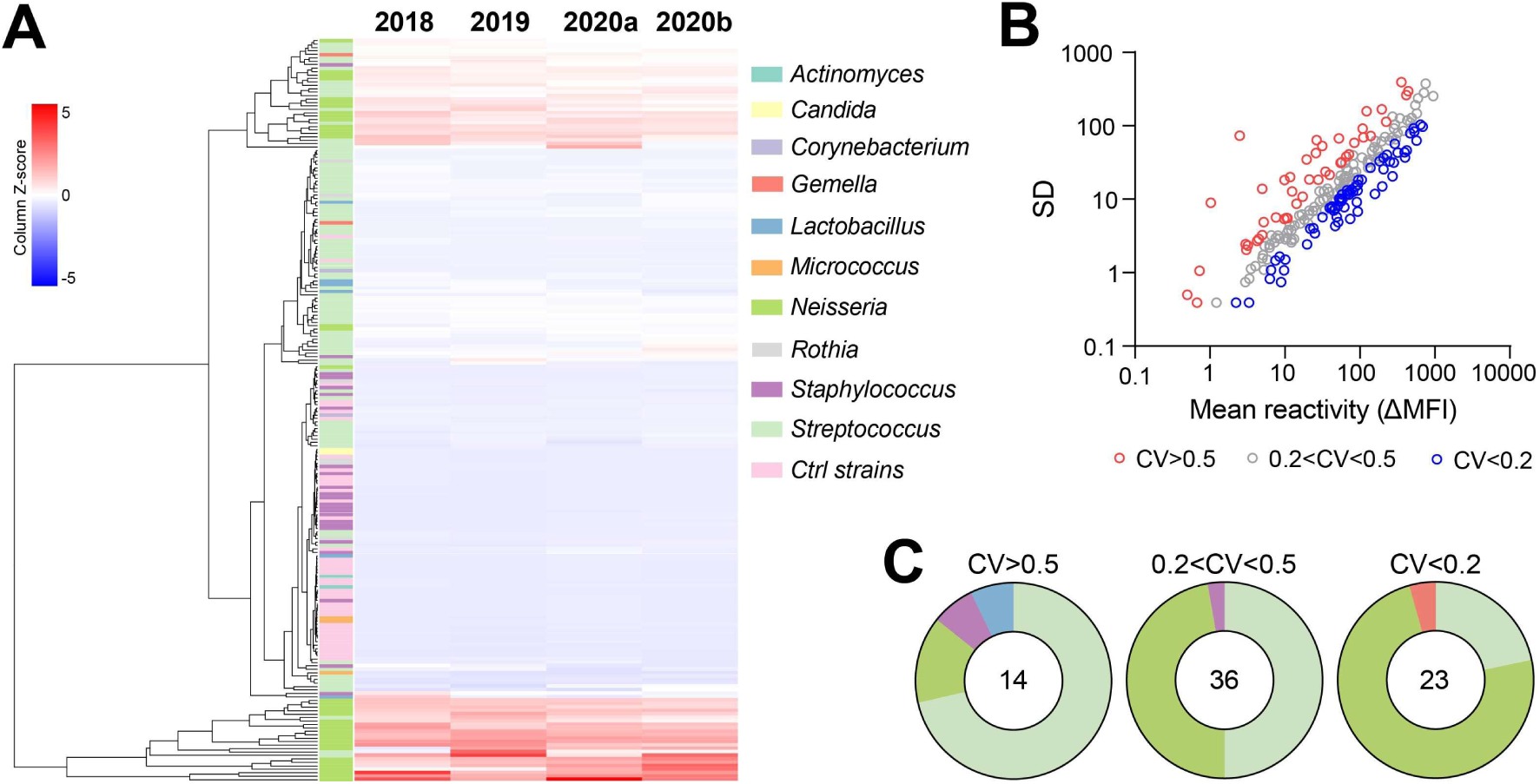
Conservation of serum IgA reactivity over time. (A) Heat map showing reactivity of serum IgA isolated from a single donor (CD1629) at four discrete time points. Reactivity was assessed against all bacterial strains (rows, n = 216) by flow cytometry. (B) Correlation between mean reactivity and SD across the four time points. For each strain, the coefficient of variation (CV) was calculated as SD divided by mean reactivity. (C) Grouping of strains according to variation in reactivity over time. Only strains with ΔMFI values >10% of the maximum signal in at least one time point are displayed. Colors indicate genera as in (A).

### Serum IgA does not reflect the reactivity of duodenal plasma cells

We next wished to address the relationship between serum IgA and mucosal IgA secreted from lamina propria plasma cells. To this end, we first assessed secretion of bacteria-binding IgA in duodenal biopsy single-cell suspensions by ELISPOT, using a selection of whole, fixed bacteria as antigens (Fig. 3A). When comparing IgA reactivity in matched serum samples and duodenal biopsies collected from a single donor at two discrete time points, we observed different reactivity patterns (Fig. 3B and 3C). The gut plasma cells did not show the same preference for *Neisseria* as we observed for serum IgA. Rather, there was broad reactivity toward most of the included strains. In addition, whereas serum IgA reactivity was conserved over time, the reactivity toward individual strains was more variable in the gut. The fact that gut biopsies, unlike serum, only capture IgA secretion from local plasma cells might result in bigger variation between samples. However, lamina propria plasma cells are believed to arise through seeding from the blood after induction in gut-associated lymphoid tissues (GALT).^28^ As such, duodenal plasma cells are not the result of a local immune response but likely represent an average of immune responses induced in GALT. Based on our results, these immune responses appear to target a broader range of bacteria and be more dynamic than systemic IgA responses.

**Figure 3.**
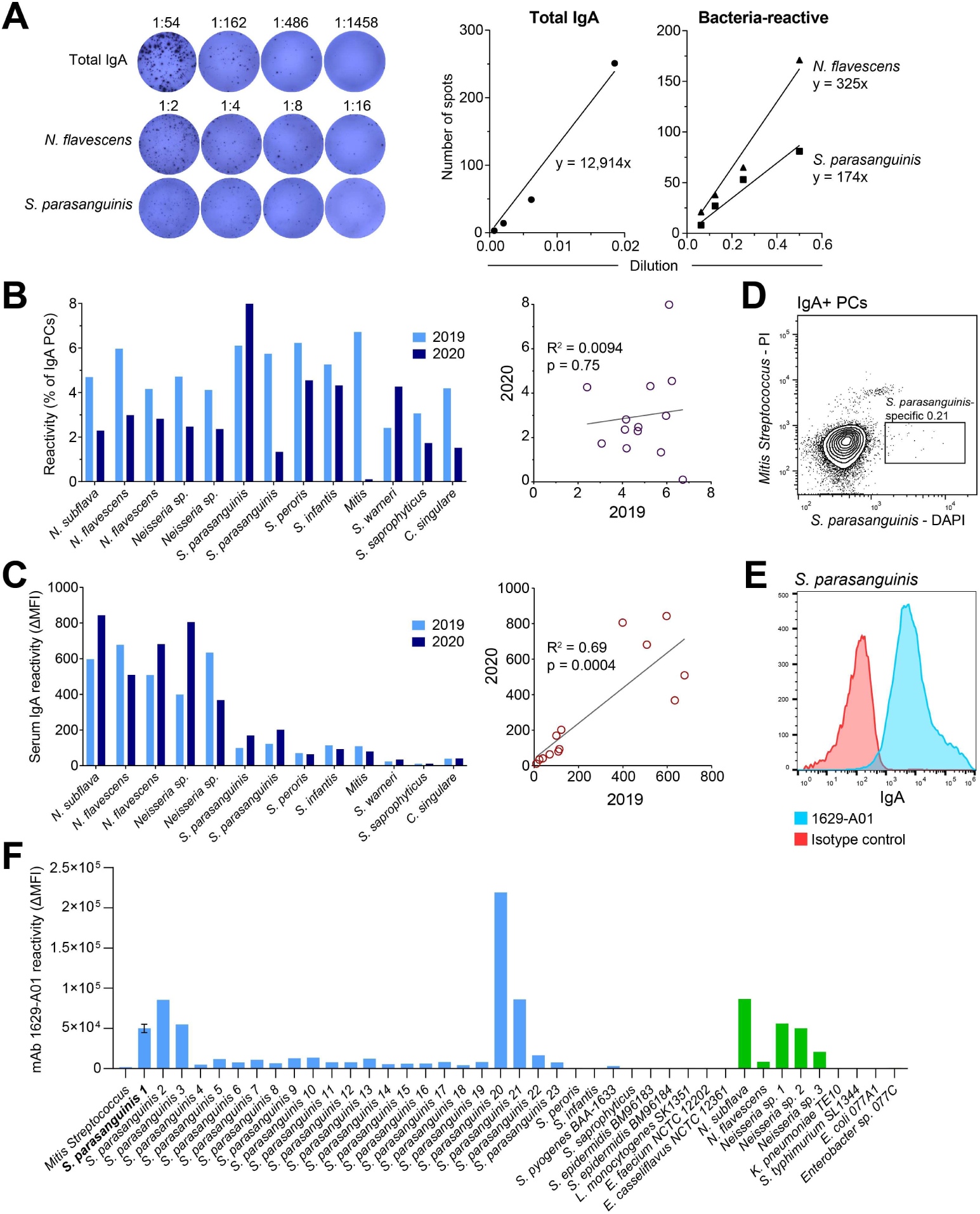
Reactivity of gut plasma cells to bacteria. (A) Detection of total and bacteria-reactive IgA plasma cells in duodenal biopsy single-cell suspensions by ELISPOT. Images and quantification of spots are shown for two representative isolates. (B and C) Comparison of plasma cell reactivity by ELISPOT (B) and serum IgA reactivity by flow cytometry (C) against individual isolates. Duodenal biopsies and serum samples were obtained from a single donor (CD1629) at two discrete time points. (D) Detection of bacteria-specific IgA plasma cells by staining of duodenal biopsy single-cell suspension with labeled isolates in flow cytometry. (E) Flow cytometry plot showing reactivity of mAb 1629-A01 generated from a single *S. parasanguinis-*binding plasma cell against the strain that was used for plasma cell isolation. (F) Reactivity of mAb 1629-A01 against a selection of isolates assessed by flow cytometry. The strain that was used for isolation of 1629-A01 is indicated in bold. Error bar indicates SD based on three experiments.

To see if we could isolate bacteria-specific plasma cells from duodenal biopsy single-cell suspensions, we selected an *S. parasanguinis* isolate that had shown consistent IgA reactivity in ELISPOT to use as DAPI-labeled antigen for plasma cell staining. In addition, we included a propidium iodide-labeled *Mitis Streptococcus* isolate as negative control antigen to increase specificity and gate out dead cells. A small population of plasma cells was observed to bind specifically to *S. parasanguinis* (Fig. 3D). After single-cell sorting, we cloned the heavy and light chain variable regions from one of the *S. parasanguinis*-binding plasma cells and expressed them recombinantly in a full-length human IgA1 format (Fig. 3E). Upon incubation with a selection of bacteria, this monoclonal antibody (mAb) showed varying degree of binding to different strains of *S. parasanguinis*. No binding was observed to other streptococci, but the mAb also recognized *Neisseria* species (Fig. 3F). This behavior is in line with the cross-species reactivity that has been observed frequently among gut plasma cells,^29^ and it suggests that the mAb binds specifically to an antigenic structure that is shared between *S. parasanguinis* and *Neisseria*. Although we have only analyzed a single mAb, it is likely that the broad IgA reactivity to bacteria that we observe in duodenal biopsies can be ascribed to widespread cross-species reactivity.

### Gut and BM plasma cells have different Ig repertoires

Since a large fraction of serum IgA is believed to be produced in the BM,^6^ we next sought to compare BM and gut plasma cells. To address the relationship between these two plasma cell populations in an antigen-specific manner, we took advantage of the prominent IgA response against the self-antigen transglutaminase 2 (TG2) in celiac disease.^30^ We have previously demonstrated a clonal relationship between TG2-binding duodenal plasma cells and purified anti-TG2 serum IgA in celiac disease patients.^31^ While TG2-specific plasma cells could readily be detected in duodenal biopsies, we were not able to identify a corresponding population in paired BM aspirates of celiac disease patients (Table S1) by flow cytometry or ELISPOT (Fig. S3). Further, droplet-based scRNA-seq in combination with CITE-seq^32^ showed that TG2-binding gut plasma cells generally did not have clonally related cells in the BM (Fig. 4A). Only a single TG2-binding clone included both gut and BM plasma cells. However, this clone also comprised non-TG2-binding cells, and it may therefore not be truly TG2 specific. Although BM plasma cells did not appear to be clonally related to TG2-specific plasma cells in the intestinal lamina propria, V(D)J sequence analysis revealed several other clonotypes spanning both compartments in all four donors analyzed (Fig. 4B). Most clones, however, were unique to either gut or BM. As previously observed, many TG2-binding gut plasma cells were negative for surface CD27.^33^ By contrast, most BM plasma cells displayed high surface CD27 levels (Fig. 4A and 4C). In agreement with the lack of correlation between serum and gut IgA against bacteria, these results indicate that strong mucosal IgA responses are not necessarily mirrored by corresponding BM plasma cell populations.

**Figure 4.**
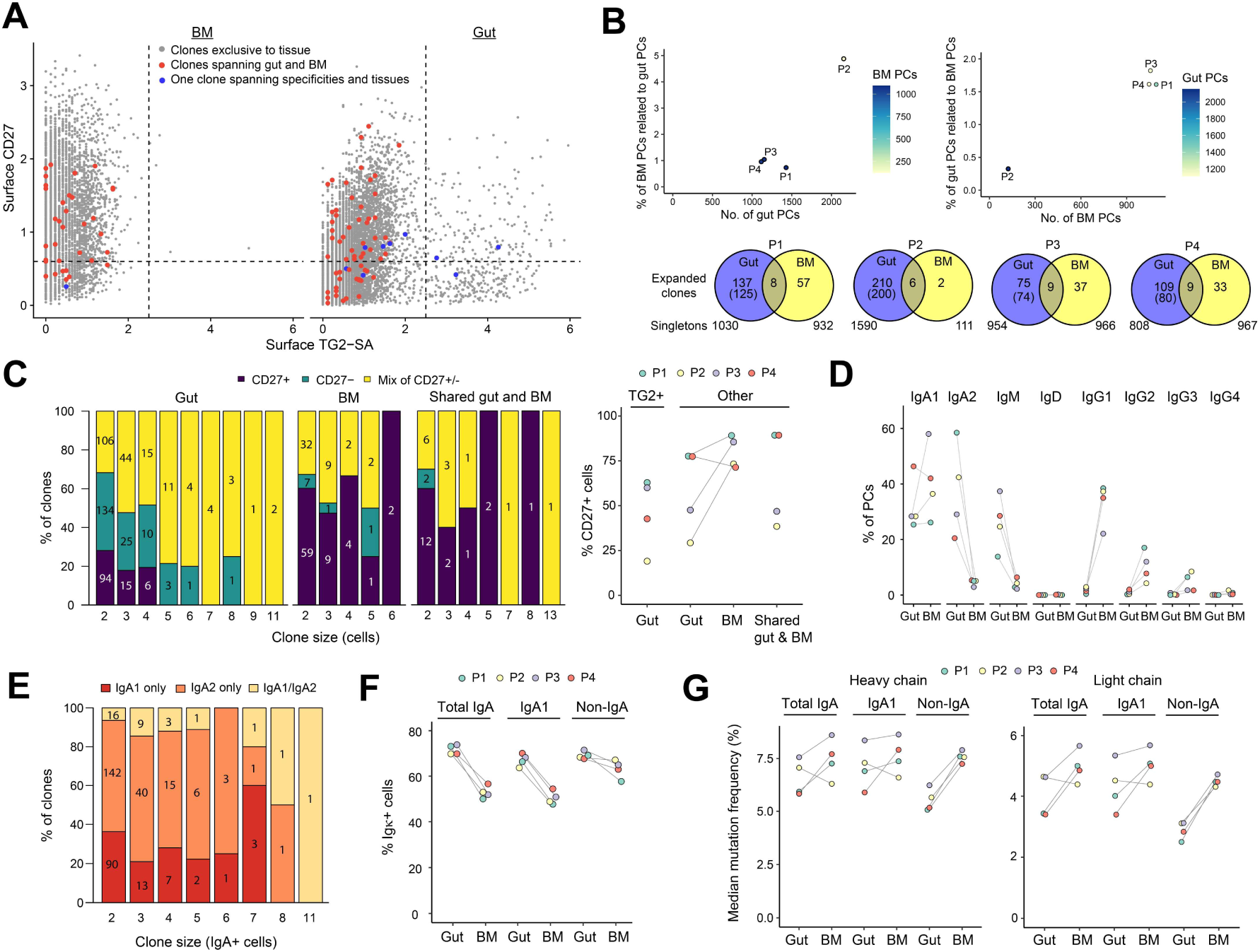
Ig sequence properties of gut and BM plasma cells. (A) Cumulated CITE-seq data showing surface CD27 expression and binding to streptavidin (SA)-coupled TG2 among BM and gut plasma cells isolated from four patients (P1-P4). (B) Clonal overlap between gut and BM plasma cells (PCs) as a function of the number of cells obtained from each compartment. The Venn diagrams indicate overlap among expanded clones comprising two or more cells. Numbers in brackets indicate non-TG2-reactive gut clonotypes. (C) Surface CD27 expression among BM and gut clonotypes or clonotypes spanning both compartments as determined by CITE-seq. Numbers within bars indicate the number of clones of a particular size that are either entirely CD27^+^, entirely CD27^−^ or containing a mix of CD27^+^ and CD27^−^ cells. Only non-TG2-binding clonotypes are included in the left panel. The right panel indicates frequency of CD27^+^ plasma cells among TG2-binding or non-TG2-binding (other) populations isolated from gut or BM or belonging to shared clonotypes of individual patients. (D) Isotype distribution among non-TG2-binding gut and BM plasma cells in individual patients. (E) Subclass expression among expanded IgA^+^ clonotypes in the gut. Numbers within bars indicate the number of clones of a particular size that are either restricted to IgA1, restricted to IgA2 or containing a mix of IgA1- and IgA2-expressing cells. (F) Kappa light chain usage among plasma cells with the indicated isotypes isolated from gut or BM of individual patients. In the gut, non-IgA cells are mostly IgM^+^, whereas BM non-IgA cells are mostly IgG^+^. (G) V-gene mutation levels in heavy and light chains of plasma cells with the indicated isotypes isolated from gut or BM of individual patients.

As expected, the majority (on average 70%) of all plasma cells in the gut expressed IgA, and almost all the remaining cells expressed IgM (Fig. 4D). In the BM, there was approximately equal levels of IgA- and IgG-expressing plasma cells and very few IgM-expressing cells. IgG expression was biased toward the IgG1 subclass followed by IgG2 and IgG3, reflecting the distribution of serum IgG.^34^ As observed in serum,^35^ IgA production in the BM was almost exclusively of the IgA1 subclass, whereas the duodenal plasma cell population showed approximately equal levels of IgA1 and IgA2. Interestingly, most of the expanded gut clonotypes were restricted to either IgA1 or IgA2, and relatively few clones contained both subclasses (Fig. 4E).

The striking difference in subclass distribution between duodenal and BM plasma cells suggests that the two populations largely comprise different IgA repertoires. This notion was confirmed by assessing heavy and light chain V- and J-gene usage (Fig. S4A-S4C). Compared with the gut, BM showed a lower frequency of plasma cells using kappa light chains (Fig. 4F).

Since gut IgA1-expressing cells tend to use lambda light chains more often than IgA2-expressing cells, this difference could be related to the difference in subclass distribution between BM and gut. However, the lower kappa/lambda ratio in BM was also clear when only considering IgA1-expressing cells. This observation may suggest that BM plasma cells have undergone more secondary rearrangements than gut plasma cells, leading to increased switching from kappa to lambda light chains.^36^ In support of extensive Ig diversification among BM plasma cells, we observed that IgA^+^ cells from BM harbored more heavy and light chain mutations than IgA^+^ cells from gut in three out of four patients when considering all plasma cells (Fig. 4G) and in all four patients when only considering surface CD27^+^ cells (Fig. S4D).

### Distinct transcriptomic profiles of BM and gut plasma cells

To further characterize potential differences between BM and gut plasma cells, we analyzed the transcriptomic profile of the isolated cells. Dimensionality reduction showed that plasma cells clustered according to their tissue of origin with IgA cells being interspersed between IgG and IgM cells in the BM and gut clusters, respectively (Fig. 5A). Accordingly, a total of 436 differentially expressed genes (DEGs) were identified when comparing total BM and gut plasma cells, while IgA and non-IgA cells had similar gene expression profiles within each compartment (Fig. 5B and 5C). DEGs included known regulators of lymphocyte migration and persistence in the BM (*CXCR4, KLF2*) and gut (*CCR10, RGS1*).^37–41^ BM plasma cells also expressed higher levels of the TNF family receptor TACI (*TNFRSF13B*), which recognizes the key cytokines B-cell activating factor (BAFF) and a proliferation-inducing ligand (APRIL). In particular APRIL is believed to be critical for maintenance of plasma cells in BM survival niches.^37^

**Figure 5.**
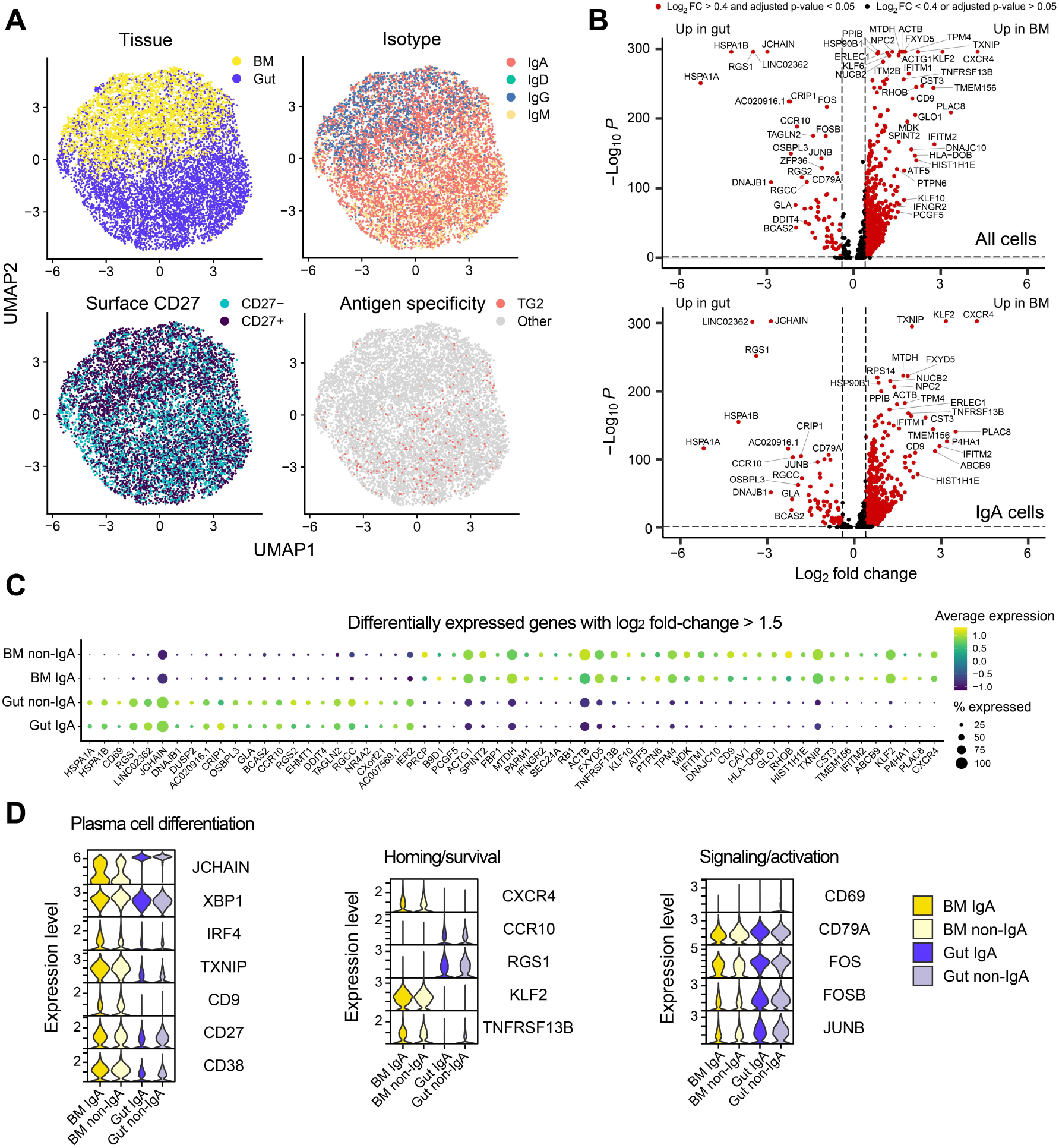
Differential gene expression between gut and BM plasma cells. (A) Uniform manifold approximation and projection (UMAP) plot based on scRNA-seq data obtained from plasma cells isolated from BM or gut of four patients. Surface CD27 expression and TG2 binding were determined by CITE-seq. (B and C) Differential gene expression between gut and BM plasma cells. The cells were subdivided according to isotype as indicated. BM non-IgA cells are primarily IgG^+^, whereas gut non-IgA cells are primarily IgM^+^. (D) Expression of selected genes associated with plasma cell differentiation, homing/survival and signaling/activation in the indicated populations.

Compared to the cells isolated from gut biopsies, BM plasma cells showed higher expression of genes typically associated with plasma cell differentiation, including the transcription factors *XBP1* and *IRF4*, the surface markers *CD9* and *CD38*, and the ER stress-related factor *TXNIP* (Fig. 5D).^37,42,43^ In agreement with the elevated frequency of surface CD27^+^ cells (Fig. 5A), BM plasma cells also showed higher expression of *CD27* at the transcriptional level (Fig. 5D). Gut plasma cells, on the other hand, expressed higher levels of genes associated with activation and B-cell receptor (BCR) signaling (*CD69*, *CD79A, FOS, JUNB*).^44,45^ These results may indicate that BM plasma cells are more terminally differentiated, while gut plasma cells could be involved in active sensing of mucosal antigens via surface IgA and IgM BCRs.^46^

In agreement with J chain being considered a common marker of plasma cell differentiation,^47^ we observed *JCHAIN* to be expressed across isotypes both in BM and gut plasma cells (Fig. 5B-5D). However, the expression level was significantly higher in the gut compared to the BM. The amount of produced J chain likely controls IgA dimerization, thus explaining why BM plasma cells would mainly secrete monomeric IgA, while gut plasma cells would secrete dimeric IgA.

### Identification of bacteria-specific plasma cell clonotypes spanning gut and BM

The plasma cell clonotypes that span gut and BM were found to have an isotype distribution similar to that of the general plasma cell population in the duodenum (Fig. 6A-C). Thus, most of the BM cells with clonally related cells in the gut expressed IgA with an approximately equal distribution between IgA1 and IgA2. To characterize the reactivity of these clones, we expressed ten of the BM-derived V(D)J sequences as full-length human IgA1 mAbs (Table S4). As expected, the mAbs did not bind recombinant TG2, nor did they display polyreactivity when tested against a panel of model antigens (Fig. S5). However, one mAb (2046-5BM) bound DNA, and another mAb (558-6BM) showed reactivity to lipopolysaccharide (LPS) of *Escherichia coli* strain O55:B5. Importantly, 558-6BM did not bind to LPS of *E. coli* strain O111:B4 or LPS from *Klebsiella pneumoniae*, nor did it bind ultrapure LPS from O55:B5 (Fig. 6D). Binding was slightly increased upon treatment of the standard O55:B5 LPS with the peptidoglycan-cleaving enzyme mutanolysin, whereas treatment with proteinase K led to complete loss of reactivity. We therefore conclude that mAb 558-6BM recognizes a protein contaminant present in the standard O55:B5 LPS preparation. Western blot analysis revealed that this contaminant is a small ∼10 kDa protein (Fig. 6E). In addition, 558-6BM showed binding to a high-MW component that disappeared upon mutanolysin treatment, suggesting that the antigen can be attached to high-MW peptidoglycan. The only *E. coli* protein known to form covalent links to peptidoglycan is major outer membrane lipoprotein (Lpp, also known as Braun’s lipoprotein), which is a 58 aa peptide attached to the outer membrane via three acyl groups.^48^ Importantly, 558-6BM showed binding to the recombinant peptide part of Lpp albeit with reduced affinity compared to O55:B5 LPS (Fig. 6F), suggesting that the mAb recognizes the combined lipid and protein constituents of Lpp.

**Figure 6.**
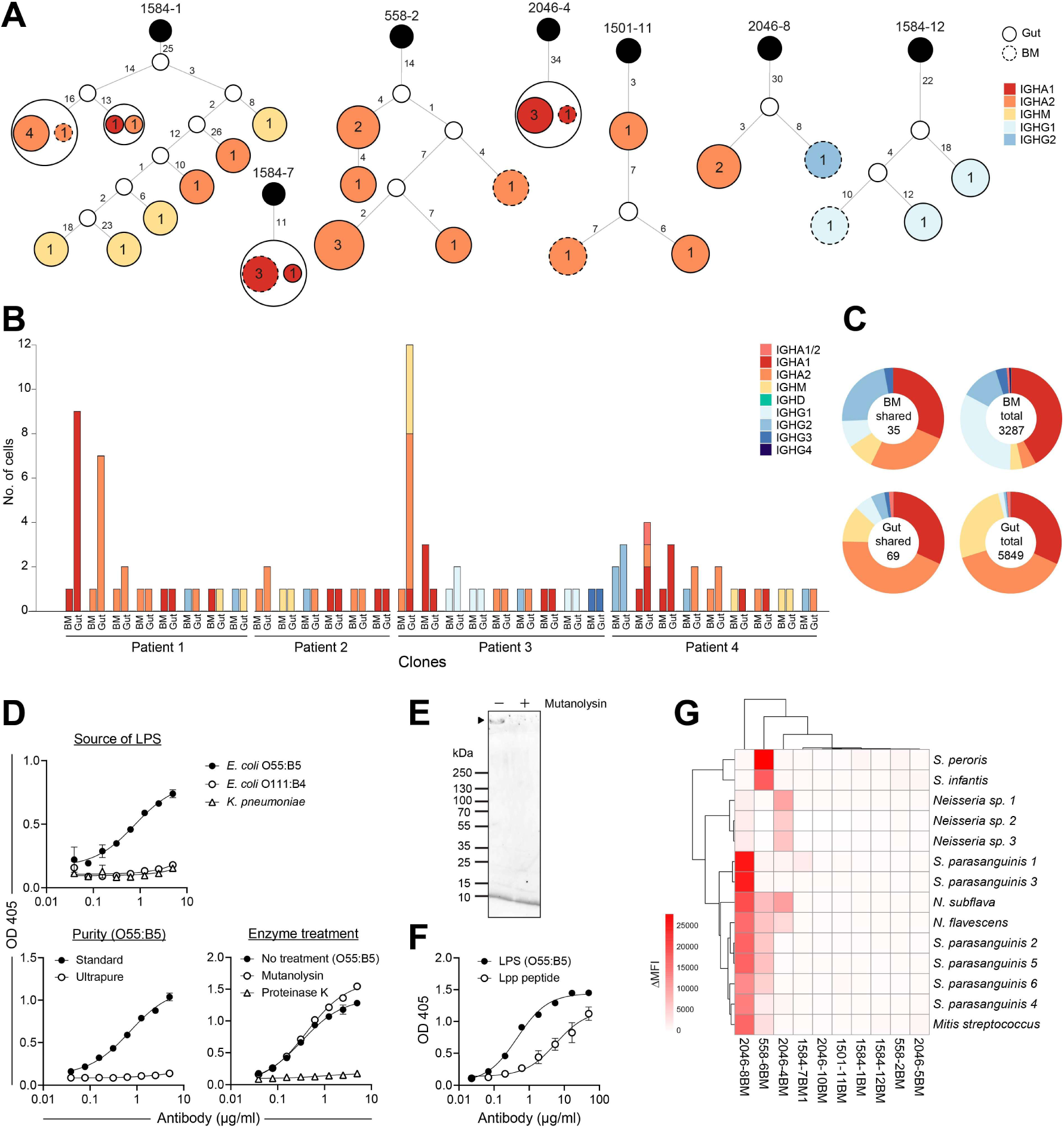
Reactivity of plasma cell clonotypes spanning gut and BM. (A) Examples of lineage trees showing plasma cell clonotypes spanning gut and BM. Observed sequences are colored according to isotype. Inferred sequences are indicated as smaller white circles, and predicted germline sequences are black. Circle sizes indicate number of cells with identical Ig sequence, and numbers next to edges indicate mutations (nt). (B and C) Overview of Ig isotype usage and number of cells in clonotypes spanning gut and BM in four celiac disease patients. Each clone is represented by one BM bar and one gut bar in (B). Isotype usage among all cells belonging to shared clones is compared to the total populations of BM and gut plasma cells in (C). IgA^+^ cells that could not be assigned to a specific subclass are indicated as IGHA1/2. IgG^+^ cells that could not be assigned to a specific subclass were excluded (in total 34 cells in BM and 1 cell in gut). (D) ELISA binding curves showing reactivity of mAb 558-6BM with LPS of different bacterial sources, purity, and after enzymatic treatment. Error bars represent range of sample duplicates. (E) Western blot showing reactivity of mAb 558-6BM with components of *E. coli* O55:B5 LPS before and after treatment with mutanolysin. In addition to a band at ∼10 kDa, a high-molecular weight band (indicated with black triangle) is present in the sample without mutanolysin. (F) Comparison of mAb 558-6BM reactivity against *E. coli* O55:B5 LPS and *E. coli* Lpp (peptide component) by ELISA. Error bars represent range of sample duplicates. (G) Reactivity of ten mAbs generated from clonotypes spanning gut and BM with a selection of bacterial isolates as determined by flow cytometry.

To test if the generated mAbs would be reactive with bacteria of the upper GI tract, we incubated them with selected strains from our panel of bacterial isolates (Fig. 6G). Three mAbs, including 558-6BM, bound selectively to some of the strains. Two mAbs (2046-8BM and 558-6BM) displayed cross-species reactivity, as they bound to several phylogenetically distant species. The third mAb (2046-4BM), however, showed specificity to *Neisseria* species. This mAb thus reflects the observed reactivity of polyclonal serum IgA. Unlike the other two bacteria-reactive mAbs, 2046-4BM was generated from an IgA1-expressing plasma cell (Fig. 6A and Table S4), further suggesting that it could be more representative of serum IgA antibodies. With three out of ten mAbs showing reactivity to a limited collection of isolates, our results suggest that plasma cells secreting antibodies to bacteria of the upper GI tract are common in the BM.

### Bacteria of the upper GI tract express IgA-modifying enzymes

The data presented so far were obtained from biological samples of patients with confirmed or suspected celiac disease. To test if reactivity toward *Neisseria* is a general feature of systemic IgA, we purified serum IgA from 13 individuals comprising both celiac disease patients and healthy donors and assessed binding to selected bacterial strains (Fig. 7A). Except for one donor, who had high IgA levels to *Rothia mucilaginosa*, all donors showed highest IgA reactivity to the included *Neisseria* species (*N. subflava*). These results thus confirm that *Neisseria* species are commonly targeted by systemic IgA across individuals in a celiac disease-independent manner.

**Figure 7.**
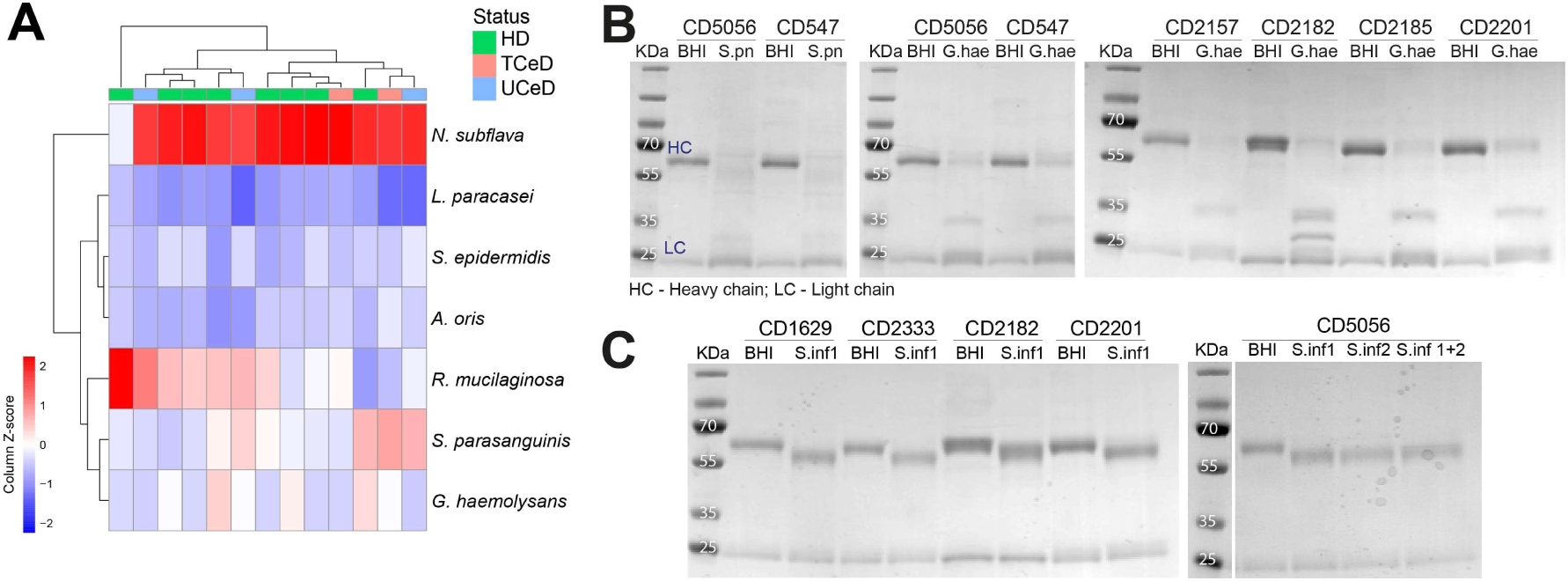
IgA-modifying activity of secreted bacterial enzymes. (A) Heat map showing IgA reactivity against a selection of bacterial isolates as determined by flow cytometry. Total IgA was purified from serum of healthy donors (HD), untreated celiac disease patients (UCeD) or celiac disease patients treated with a gluten-free diet for at least one year (TCeD) (B) Assessment of IgA integrity by SDS-PAGE after incubation with bacterial culture supernatants. Brain Heart Infusion (BHI) broth alone was used as negative control and supernatant of *Streptococcus pneumoniae* (S.pn), which is known to express IgA1 protease, was used as positive control. One isolate of *Gemella haemolysans* (G.hae) was found to have IgA-degrading activity. The effect was observed using purified serum IgA of six individual donors. (C) Detection of IgA heavy chain mobility shift after incubation of purified serum IgA of five donors with supernatant of two isolates of *Streptococcus infantis* (S.inf) used either alone or in combination.

Selective generation of systemic IgA against certain bacteria can potentially be explained by the ability of other bacteria to avoid such responses. One way in which bacteria may escape recognition by IgA-expressing B cells is via secretion of proteases that specifically cleave IgA. Proteases specific for IgA1 are expressed by pathogenic *Neisseria* species (*N. meningitidis* and *N. gonorrhea*) but not by commensal *Neisseria* such as the ones described in this study.^49^ Among the second large group of bacteria in the upper GI tract, the streptococci, the pathogen *S. pneumoniae* is known to express IgA1 protease.^50–52^ In addition, some non-pathogenic streptococci have been found to encode IgA1 proteases in a strain-specific manner.^53–55^ To test if the bacteria we isolated from duodenal biopsies express IgA-degrading enzymes, we incubated purified serum IgA with supernatant of bacterial cultures and assessed protein integrity by SDS-PAGE. Of 30 isolates tested (17 streptococci, 3 staphylococci, 2 isolates each of *Neisseria*, *Gemella* and *Corynebacterium*; 1 isolate each of *Rothia*, *Actinomyces*, *Lactobacillus* and *Candida*), we found that one (*Gemella haemolysans*) was able to degrade IgA (Fig. 7B), in agreement with a previous observation.^56^ In addition, we observed a small shift in the molecular weight of the IgA heavy chain upon incubation with two different isolates of *Streptococcus infantis* (Fig. 7C). This shift can most likely be explained by bacterial glycosidases, which cleave off IgA-associated glycans, in agreement with previous observations.^54,57^ These results demonstrate that bacteria in the upper GI tract can express IgA-modifying enzymes, which might play a role in shaping IgA responses against bacteria. Importantly, the observed activities were specific to IgA, as we did not see any modification of purified serum IgG (Fig. S6).

## DISCUSSION

The largest population of plasma cells in the human body is found in the intestinal lamina propria. Many of these plasma cells secrete antibodies that specifically bind to bacterial antigens, leading to efficient coating of the intestinal microbiota.^10,12,58,59^ The antibodies show signs of antigen-driven selection, and they are believed to result from B-cell activation in GALT, where Peyer’s patches are considered particularly important for induction of gut IgA responses.^28^ Accordingly, a clonal relationship has been demonstrated between B cells in Peyer’s patches and lamina propria plasma cells.^60^ Generation of mAbs from gut plasma cells has demonstrated that many clones are able to recognize several phylogenetically unrelated bacteria, a phenomenon termed cross-species reactivity.^29,59^ This behavior is likely the result of B-cell responses being directed against surface structures, particularly glycans, which can be shared across many bacterial species.^12,61^ In line with previous reports, we here show that duodenal IgA-secreting plasma cells display broad reactivity toward bacteria isolated from duodenal biopsies. Further, by staining plasma cells with a strain of *S. parasanguinis*, we obtained a mAb that showed cross-reactivity with *Neisseria* species. Thus, even when using a single bacterium to target antigen-specific plasma cells, the identified cells display cross-species reactivity.

The broad reactivity of gut plasma cells toward bacteria was not reflected by serum IgA. Instead, we observed a striking selection for certain bacteria that are commonly present in the upper GI tract. Typical species found in the colon, such as *E. coli* and *Enterococcus faecium*, on the other hand, did not show high serum IgA reactivity. Among isolates from the upper GI tract, serum IgA showed preferential binding to *Neisseria* species, followed by streptococci. This reactivity pattern was found both in celiac disease patients and in healthy donors. The response was stable over time and did not appear to distinguish between different species within the *Neisseria* genus. Our results with polyclonal serum IgA were corroborated by generation of a *Neisseria*-specific mAb from an IgA1 plasma cell clonotype that spanned BM and gut. Taken together, systemic IgA antibodies against the microbiota seem to be biased toward binding of *Neisseria*, but do not show specificity down to the strain or species level.

Although some strains were associated with private antibody responses, individual donors generally showed serum IgA reactivity not only to their own bacterial isolates, but also to isolates of other individuals. Preferential recognition of “self” bacteria by serum IgA was demonstrated in a recent mouse study.^17^ In that study, however, mice were colonized with a limited set of defined bacterial strains. In humans, on the other hand, the diversity of the microbiota will likely drive selection for antibodies with broader reactivity, since B cells that can recognize multiple bacterial species will have an advantage in competition for T-cell help.^62^

By conducting scRNA-seq on paired samples of duodenal biopsies and BM aspirates, we demonstrate differences in plasma cell V(D)J repertoire and transcriptomic profile between the two compartments. Compared with gut plasma cells, BM IgA^+^ plasma cells had lower expression of J chain and a strong bias toward the IgA1 subclass, consistent with them being a major source of monomeric serum IgA. Among secondary lymphoid tissues, a similar low frequency of IgA2-expressing B cells has only been demonstrated in spleen, peripheral lymph nodes and tonsils.^18,19^ Peyer’s patches and mesenteric lymph nodes, on the other hand, display higher IgA2 levels.^63^ The question whether IgA2 mainly arises via direct switching from IgM or via sequential switching from IgA1 (or other isotypes) is debated.^64–67^ Potentially, different segments of the gut could be governed by different class switching mechanisms, as the small and large intestine harbor clonally distinct plasma cell populations.^66,68^ Our analysis of duodenal plasma cells indicates that IgA1 and IgA2 cells largely belong to different clonotypes, suggesting that the IgA1/IgA2 distribution is controlled by primary B-cell activation in secondary lymphoid organs. These observations lead us to propose that IgA-expressing plasma cells in BM and gut originate from immune responses at discrete anatomical sites. Whereas lamina propria plasma cells are thought to derive mainly from Peyer’s patches, IgA-expressing BM plasma cells are likely generated in another mucosa-associated lymphoid tissue. Waldeyer’s ring, including the palatine tonsils and adenoid, is a group of lymphoid tissues in the naso- and oropharynx that may be particularly important for generation of systemic IgA. Like Peyer’s patches, these structures are heavily exposed to antigens and a site of continuous immune cell activation, germinal center formation and IgA class-switching.^69,70^ Interestingly, a tonsil-derived activated B-cell subset was shown to have homing properties distinct from those of GALT-derived B cells and was also demonstrated to populate the BM.^71^ Compartmentalization of mucosal immune responses is also supported by the observation that oral and nasal immunization give rise to different types of antibody responses with distinct mucosal distributions.^72,73^ Further, in agreement with our observation of pronounced systemic IgA reactivity toward *Neisseria* species, commensal *Neisseria* have been shown to make up a large fraction of bacteria in tonsil swabs.^74^ It is possible that the *Neisseria* isolates we obtained from duodenal biopsies were picked up from the pharynx or oral cavity through the endoscopy procedure.

Despite the different anatomical origins of BM and gut plasma cells, we identified clonotypes spanning both compartments. Interestingly, BM plasma cells belonging to such shared clonotypes had an IgA1/IgA2 distribution mirroring that of the duodenum with more IgA2-expressing cells than seen in the general BM population. The inductive site for these shared clonotypes is therefore likely GALT. It is possible that initial activation in GALT leads to formation of circulating memory cells that can become re-activated and differentiate into BM-homing plasma cells upon encounter of the same antigen in another site, such as tonsils. In agreement with this idea, two mAbs generated from IgA2 or IgA2/IgG2 clonotypes spanning gut and BM displayed broad reactivity to bacteria and could thus have been activated against different species populating distinct mucosal sites. Notably, one of these mAbs (558-6BM) bound Lpp of *E. coli*, but at the same time recognized surface antigen on gram-positive bacteria. Importantly, lipoproteins are found in all bacteria,^75^ and it is conceivable that 558-6BM reacts with Lpp homologs present in phylogenetically distant species.

To address the relationship between antigen-specific mucosal and systemic IgA, we took advantage of the autoantibody response against TG2 in celiac disease patients. Although TG2-specific plasma cells were abundant in patient duodenal biopsies, we were not able to identify similar cells in paired BM aspirates. We have recently shown that TG2-specific IgA^+^ B cells in blood samples of celiac disease patients have an activated pre-plasma cell phenotype and are homing to the small intestinal mucosa.^33^ Interestingly, most of the TG2-specific cells, both circulating B cells and lamina propria plasma cells, were negative for the classical memory marker CD27 and harbored low levels of somatic V-gene mutations. Since both somatic hypermutation and expression of CD27 is induced in germinal center B cells,^76,77^ it is plausible that TG2-specific memory B cells and plasma cells are generated through an extrafollicular activation route. By contrast, we here show that BM plasma cells generally showed high expression of CD27 and were highly mutated, consistent with the notion that most long-lived, BM-residing plasma cells are derived from germinal center reactions.^78^ The fact that formation of anti-TG2 serum IgA is a short-lived antibody response that is completely dependent on dietary gluten^79^ also points to a lack of TG2-specific plasma cells situated in BM survival niches. In active celiac disease, the source of anti-TG2 serum IgA is therefore likely short-lived plasma cells located in secondary lymphoid tissues or circulating plasmablasts.^33^

Given the diversity of the microbiota populating mucosal surfaces, the observation that systemic IgA against bacteria preferentially bound to *Neisseria* species was surprising. To explain the selective response against *Neisseria*, we assessed whether our isolates expressed IgA-degrading enzymes. Potentially, a bacterium secreting IgA-specific protease would be able to cleave the BCR of nearby IgA^+^ B cells. It would thereby escape recognition, prevent B-cell activation and, thus, avoid formation of specific anti-bacterium antibodies. Among the tested strains, we identified a single isolate of *G. haemolysans* which specifically cleaved purified serum IgA. In addition, we identified two isolates of *S. infantis* which displayed IgA-specific glycosidase activity. Glycosylation of the IgA heavy chain constant region has been shown to mediate non-canonical binding to some bacteria via glycan-glycan interactions.^3,80^ Thus, cleavage of such glycans by bacterial glycosidases will likely modulate IgA binding to some species. Although we could only demonstrate secretion of IgA-modifying enzymes in a few isolates grown *in vitro*, it is possible that such activities will be more widespread during bacterial colonization of the human body. Bacterial IgA-modifying enzymes could thereby contribute to shaping of the IgA response against the microbiota. Since we observed IgA-modifying activities in species of *Gemella* and *Streptococcus*, it is tempting to speculate that they are part of the explanation for the overall low serum IgA reactivity against those genera.

Taken together, our study demonstrates selective generation of systemic IgA against certain bacteria in the upper GI tract. These antibody responses appear to depend on inductive sites that are distinct from those of lamina propria plasma cells in the gut. Thus, lymphoid tissues of the upper aerodigestive tract might be particularly important for generation of BM-residing plasma cells that are responsible for stable production of serum IgA. Distinct formation of secretory and systemic IgA has implications for development of mucosal vaccines. Hence, if both systemic and gut mucosal protection is desired, it might be necessary to target tonsils as well as GALT.

### Limitations of the study

To study antibody responses against bacteria of the upper GI tract, we isolated strains from duodenal biopsies of individual donors. The bacteria obtained by this approach do not represent the complete microbiome. Rather, our study is limited to the bacteria present in the samples and those that could be cultured *in vitro* using standard culture media and aerobic conditions. In addition, some bacteria might change their phenotype or display phase variation in culture, causing them to have different antigenic properties than they would have in the human body. The use of whole, fixed bacteria for assessing antibody reactivity limits our study to surface-exposed antigens. While we might lose detection of some antibodies, it has been demonstrated that most bacteria-reactive mucosal plasma cells are specific to cell wall and membrane-associated antigens.^81,82^

Based on differences in molecular composition and antigen reactivity, we hypothesize that systemic and secretory IgA responses are induced at discrete anatomical sites and that lymphoid tissues of the upper aerodigestive tract are important for generation of BM-residing IgA^+^ plasma cells. To obtain formal proof for this model, it will be necessary to assess the degree of clonal overlap between B-cell populations in tonsils and BM. Thus, future studies should aim to address the relationship between B cells/plasma cells in these two compartments.

## ACKNOWLEDGMENTS

We are thankful to all patients who donated biological material and to the staff at the Endoscopy Unit at Oslo University Hospital for help with sample collection. Marie Noer is thanked for help with identification of bacteria. In addition, we thank the following colleagues for kindly gifting us bacterial strains that helped us to develop this study: Daniel Walker, Donal Wall and Gillian Douce (University of Glasgow), José R. Penades (University of Glasgow and Imperial College London), and Tim Foster (Trinity College Dublin, University of Dublin). Flow cytometry and cell sorting experiments were conducted at the Flow Cytometry Core Facility, Oslo University Hospital, and next generation sequencing was performed at the Norwegian Sequencing Centre, University of Oslo. Raw sequencing data were stored and processed on the Tjenester for Sensitive Data (TSD) facilities, owned by the University of Oslo. This work was supported by grants from Stiftelsen KG Jebsen (project SKGJ-MED-017), the South-Eastern Norway Regional Health Authority (project 2022071) and the University of Oslo through the Scientia Fellows II program and the World-leading research program on human immunology (WL-IMMUNOLOGY).

## AUTHOR CONTRIBUTIONS

Conceptualization, F.V. and R.I.; Methodology, F.V., I.L., and R.I.; Investigation, F.V., I.L., H.H.H., C.S., A.K.S., and R.I.; Resources, G.E.T., and K.E.A.L.; Writing – Original Draft, F.V., and R.I.; Writing – Review & Editing, all authors; Supervision, J.V.B., L.M.S., and R.I.; Funding Acquisition, F.V., L.M.S., and R.I.

## DECLARATION OF INTERESTS

The authors declare no competing interests.

## MATERIALS AND METHODS

### Collection of biological samples

Blood, duodenal biopsies and BM aspirates were collected from adult patients and control subjects who had given informed consent. The diagnosis of celiac disease was made according to the guidelines of the European Society for the Study of Coeliac Disease.^83^ Ethical approval was given by the Regional Ethics Committee of South-Eastern Norway (REK ID 6544).

### Isolation of human cells

Biopsy specimens were disrupted by incubation with 1 mg/mL of collagenase (Sigma) in 2% (v/v) FBS/PBS under rotation for 1 h at 37°C. The digested samples were homogenized with a syringe and filtered through a 40 μm cell strainer. BM aspirates collected from the iliac crest were supplemented with 2 mM EDTA, before they were passed through a 40 µm cell strainer. The BM samples were diluted 1:1 with RPMI-1640, and mononuclear cells were isolated by density gradient centrifugation. All cells were cryopreserved in 50% FBS/RPMI-1640 supplemented with 10% (v/v) DMSO and stored in liquid nitrogen.

### Isolation of bacteria from duodenal biopsies

Duodenal biopsies that were used to grow bacteria were collected directly into a tube with sterile PBS. Biopsy specimens (3-4 pieces of ca. 1 mm^3^ each) and 5 mL of the PBS solution were transferred to a 25 cm^2^ vented flask and supplemented with 5 mL of 2xBMYE (2x buffered medium: 65 mM Na2HPO4, 46 mM NaH_2_PO_4_, 147 mM NaCl supplemented with 0.2% [w/v] tryptone and 0.5% [w/v] Bacto Yeast Extract [YE, Gibco]). After 24 h incubation at 37°C, 5% CO_2_, the culture media were serially diluted in 1:1 2xBMYE:PBS and plated on different agar media: Brain Heart Infusion (BHI, Oxoid) supplemented with 5% YE and 10% (v/v) defibrinated sheep blood (Thermo Fisher Scientific), Luria-Bertani agar, 2xBMYE agar, or BHI supplemented with 0.3% YE, 1.2% (w/v) casein peptone and 5% defibrinated sheep blood. The plates were incubated at 37°C, 5% CO_2_ and checked daily until day 7. Colonies were streaked after 48 h. After 3-6 passages from single colony, the strains were grown in broth for preservation as glycerol stocks. For identification, the isolates were grown on agar plates and subjected to matrix-assisted laser desorption/ionization-time of flight (MALDI-TOF) mass spectrometry using a MALDI Biotyper instrument (Bruker Daltonics). Briefly, cells were picked from a confluent growth area with a 1 µL loop and directly spotted onto a MALDI target plate in duplicate. The cells were lysed with formic acid and coated with HCCA matrix. Table S2 shows the isolated strains used in this study.

### Growth of bacterial strains

Bacteria of the *Enterobacteriaceae* family were grown on Luria-Bertani agar/broth at 37°C; *Bifidobacterium animalis* was grown in Columbia agar (home-made recipe for 1 L: 12 g tryptone, 3 g YE, 5 g BHI, 3 g Lab-lemco powder [Oxoid], 5 g NaCl, 15 g Agar), or 58 DSMZ broth (recipe for 400 mL: 4 g tryptone, 2 g YE, 2 g Bacto Soytone [Gibco], 2 g Lab-lemco powder, 4 g glucose, 0.8 g K_2_HPO_4_, 0.08 g MgSO_4_·7H_2_O, 0.02 g MnSO_4_·H_2_O, 400 µL Tween-80, 2 g NaCl, 0.02 g cysteine·HCl, 16 mL salt solution [recipe for 400 mL: 0.1 g CaCl_2_·H_2_O, 0.2 g MgSO_4_·7H_2_O, 0.4 g K_2_HPO_4_, 0.4 g KH_2_PO_4_, 4 g NaHCO_3_, 0.8 g NaCl]). All other strains were grown in agar or broth of BHI supplemented with 5% YE. Strict anaerobes were incubated at 37°C in anaerobic jars with AnaeroGen sachets (Oxoid) for 48 h. All other bacteria were grown at 37°C, 5% CO_2_ without shaking. For detection of reactivity with antibodies or gut plasma cells, bacterial strains were grown in 10 mL of broth for 24-48 h, depending on the strain, until the culture reached stationary phase. For each test, cells from 1 mL of culture were harvested, washed once with 700 µL PBS and fixed in 70 µL alcohol-free 4% formaldehyde fixative solution (Thermo Fisher Scientific). The fixed cells were washed twice with PBS and resuspended in 200 µL 1% (w/v) BSA/PBS. They were then stored at 4°C until they were analyzed. Tables S2 and S3 present all strains used in this study.

### Purification of serum IgA and IgG

Serum (1 mL) was diluted ten times with PBS and passed through a 0.2 µm filter, before total IgA was purified on a Tricorn 10/20 column (Cytiva) packed with peptide M agarose (InvivoGen). After washing with at least five column volumes of PBS, bound IgA was eluted with 0.1 M Glycine-HCl, pH 2.5 followed by immediate neutralization with 1 M Tris-HCl, pH 9. The purified IgA was concentrated and buffer exchanged into PBS using Vivaspin 20 centrifugal concentrators with 30 kDa cut-offs (Sartorius). Serum IgG was purified in an equivalent way using a HiTrap protein G column (Cytiva).

### Sorting of plasma cells

Plasma cells were identified by staining gut biopsy single-cell suspensions or BM aspirates with the following combination of antibodies: Anti-human CD38-FITC (clone HB7, Thermo Fisher Scientific), anti-human IgA-APC (clone IS11-8E10, Miltenyi), anti-human IgD-PerCP/Cy5.5 (clone IA6-2, BioLegend), anti-human CD3-Brilliant Violet 510 (clone OKT3, BioLegend), anti-human CD14-Brilliant Violet 510 (clone M5E2, BioLegend). Plasma cells were identified as large CD38^Hi^, IgD^−^, CD3^−^, CD14^−^ lymphocytes, and dead cells were excluded by labeling with LIVE/DEAD fixable Aqua or Near-IR viability stains (Thermo Fisher Scientific). For detection of TG2-specific cells, recombinant human TG2 was tetramerized on PE-conjugated streptavidin (Thermo Fisher Scientific) via a biotinylated N-terminal AviTag as previously described.^33^ Biotinylated glutathione S-transferase (GST) attached to PE/Cy7-conjugated streptavidin (Thermo Fisher Scientific) was used as irrelevant control antigen to increase staining specificity. Both tetramers were added to the cell suspensions together with labeled antibodies. For detection of bacteria-specific plasma cells, duodenal biopsy single-cell suspensions were stained with labeled bacterial strains in addition to anti-human CD38-FITC and anti-human IgA-APC. Fluorochrome labeling of bacteria was performed by incubating individual fixed isolates with 2.5 µg/mL propidium iodide or DAPI for 20 min at room temperature. After extensive washing with PBS, the bacterial pellets were resuspended in 200 µL flow buffer (2% FBS/PBS), and 5 µL was added to the biopsy suspension. Plasma cell populations were analyzed and sorted using a FACSMelody instrument (BD Biosciences).

### Droplet-based scRNA-seq

BM and gut plasma cells were identified as described above with the exception that TG2 tetramers were generated with DNA-barcoded streptavidin-PE (TotalSeq-C0951, BioLegend). In addition, a barcoded anti-human CD27 antibody (TotalSeq-C0154, BioLegend) was included, and BM and gut cell suspensions were labeled separately with anti-human Hashtag 1 (TotalSeq-C0251, BioLegend) and Hashtag 2 (TotalSeq-C0252, BioLegend) antibodies, respectively. The cells were incubated with FcR blocking reagent (Miltenyi Biotec) prior to addition of antibodies and antigens. Following incubation on ice, the cells were washed, and total plasma cells were bulk-sorted into empty microtubes that had been pre-coated with 1% BSA/PBS. Hashtagged BM and gut plasma cells from each patient were combined into the same tube to reach a total of 10,000-12,000 cells. Barcoded cDNA from four patients was prepared with the 10x Genomics Chromium Controller, followed by generation of 5′ gene expression, V(D)J-enriched and cell surface protein libraries using Chromium single cell kits (v1.0) according to the instructions from the manufacturer (10x Genomics). The libraries were pooled and sequenced on a NovaSeq 6000 instrument (Illumina) generating 150 bp paired-end reads.

### scRNA-seq quality control

The gene expression, V(D)J and cell surface protein data were processed with 10x Genomics Cell Ranger v6.0.2 using the multi and aggr functions and GRCh38 version 2020-A for gene expression and GRCh38-alts-ensembl-5.0.0 for V(D)J analysis. Quality control was performed using Seurat v5.2.1^84^ in R 4.4.3 based on the following criteria: >200 detected genes, 2000– 45000 Unique Molecular Identifiers (UMIs), <10% mitochondrial genes, >10% Ig genes and a productively rearranged BCR heavy chain of known isotype reported in the ‘‘airr_rearrangement.tsv’’ files generated by Cell Ranger multi for each patient. Cells were classified as gut-derived or BM-derived based on Hashtag labeling, which was normalized by centered log-ratio (CLR) transformation, and double positive or double negative cells were discarded. In total 3320 BM-derived and 5850 gut-derived cells passed quality control. Cells were further categorized as CD27^+^, CD27^−^, TG2-specific and other specificity by barcoded anti-CD27 and TG2 tetramer labeling, normalized by CLR transformation (gating shown in Fig. 4A).

### scRNA-seq analysis

Gene expression analysis was performed using Seurat. Due to the high level of expression of Ig genes in plasma cells, Ig genes were discarded from the analysis to focus on non-Ig transcriptional differences. The gene expression matrix was then normalized, highly variable genes were identified, and the gene matrix was scaled, regressing out unwanted variation from the number of detected genes and reads, percent mitochondrial genes and patient-specific differences. UMAP plots were created based on the first six principal components. For transcriptional differences between BM- and gut-derived cells, TG2-specific cells were removed from the dataset before normalization and scaling of the data. Differentially expressed genes (DEGs) between two populations were determined with the FindMarkers function of Seurat for genes expressed in at least 25% of the cells of either group, using Wilcoxon rank-sum test and the Benjamini-Hochberg method for correcting for multiple testing. DEGs were visualized with EnhancedVolcano v1.18.0 with adjusted p value 0.05 and log2 fold-change 0.4 as cutoffs, and dotplots were drawn for the genes with the highest log2 fold-change using the DotPlot function of Seurat. Violin plots of additional genes of interest were drawn with the VlnPlot function of Seurat.

### BCR analysis and clonal assignment

Rearranged BCR sequences and clonally related cells in the V(D)J-enriched scRNA-seq libraries were identified by Cell Ranger for each patient (see above). An expanded clone was defined as a clone consisting of two or more cells, while a singleton describes a cell not found to be clonally related to any other cell from the same donor. The BCR sequences were further analyzed using IMGT/HighV-QUEST^85^ in March 2023, and productively rearranged sequences from cells passing quality control were kept. Only the most highly expressed heavy and light chain were retained in rare cases where multiple heavy or light chains were reported for a cell. Rare sequences with too low coverage of the C-region to determine subclass were manually assigned to the main isotype instead of a specific subclass and were excluded from analyses focusing on specific subclasses. Clones of particular interest (clones comprised of cells from both gut and BM) were analyzed in more detail by BraCeR^86^ to verify their clonal relatedness determined by Cell Ranger and draw lineage trees based on both heavy and light chain, using the DefineClones function of Change-O^87^ with CDR3 nucleotide distance threshold 0.2 for clonal assignment. Repertoire analysis was done using Alakazam v1.3.0,^87^ while mutational load was analyzed with Shazam v1.2.0.^87^

### Generation of monoclonal antibodies

For generation of bacteria-reactive mAb, single IgA^+^ plasma cells binding specifically to *S. parasanguinis* were sorted into 5 µl 20 mM Tris-HCl, pH 8, 1% (v/v) Nonidet P40 Substitute, 5 U murine RNase Inhibitor (New England Biolabs). The lysed cells were flash-frozen on dry ice and stored at -70°C. Heavy and light chain variable regions were amplified by a nested RT-PCR approach as previously described.^88^ After sequencing, PCR products were reamplified using gene-specific AgeI-IGKV or AgeI-IGHV forward primers^89^ in combination with BsiWI-IGKJ^89^ or BlpI-IGHA (CAGAGGCTCAGCGGGAAGACCTTG) reverse primers. The PCR products were then cloned into vectors for expression of full-length human IgA1.^90^ For expression of clonotypes spanning BM and gut, selected variable region sequences were ordered as synthetic DNA (GenScript) and subcloned into the same expression vectors. The antibodies were expressed by transient transfection of Expi293F cells followed by purification from culture supernatants in the same way as described for serum IgA.

### Assessment of bacterial IgA reactivity

For assessment of antibody reactivity, fixed bacterial cells were incubated with purified serum IgA or recombinant IgA1 mAbs at 10 µg/mL in 1% BSA/PBS for 1 h at room temperature followed by washing and staining with goat F(ab’)2 anti-human IgA-AF647 (SouthernBiotech). The cells were analyzed on an Attune NxT flow cytometer (Thermo Fisher Scientific), and ΔMFI values were calculated by subtracting the signal obtained with control mAb (TG2-specific IgA1, clone 679-14-E06)^90^ from the signals obtained with each antibody preparation.

### ELISAs

Screening of mAbs for polyreactivity was done as previously described.^91^ Briefly, ELISA plates were coated with the following antigens in PBS: 10 µg/mL double stranded DNA from calf thymus (Thermo Fisher Scientific), single-stranded DNA (prepared by heating of double-stranded DNA for 30 min at 95°C), LPS from *E. coli* strain O55:B5 (Sigma), keyhole limpet hemocyanin (Sigma), or 5 µg/mL human insulin (Sigma). Goat anti-human Ig (SouthernBiotech) coated at 2 µg/mL was used as positive control antigen. Each antigen was incubated with 1 µg/mL of IgA1 mAbs in PBS supplemented with 0.1% (v/v) Tween-20 (PBST) followed by detection with AP-conjugated goat anti-human IgA (Sigma). The previously reported polyreactive IgG1 mAb 1341-C11^91^ was included as control and was detected with AP-conjugated goat anti-human IgG (SouthernBiotech). For detection of mAb reactivity with recombinant proteins, 3 µg/mL human TG2^92^ or 2 µg/mL *E. coli* Lpp protein (TargetMol) was coated in PBS followed by incubation with IgA1 mAbs in various concentrations and detection as described above. Comparison of mAb reactivity with different LPS preparations was done as described above using standard LPS from *E. coli* O55:B5 (Sigma), ultrapure LPS from *E. coli* O55:B5 (InvivoGen), LPS from *E. coli* O111:B4 (Sigma), or LPS from *K. pneumoniae* (Sigma). In one set of experiments, the standard LPS preparation was treated with mutanolysin (Sigma) or proteinase K (Qiagen) for 1 h at 37°C prior to coating.

### Western blotting

To characterize reactivity of mAb 558-6BM, 10 µL of 1 mg/mL LPS from *E. coli* O55:B5 (Sigma) was incubated for 1 h at 37°C with or without 1 µL mutanolysin (Sigma) followed by addition of SDS sample buffer and loading on a 4-20% Tris-Glycine gel (Thermo Fisher scientific). The samples were subjected to SDS-PAGE and transferred to nitrocellulose by semi-dry transfer. The membrane was then blocked in PBS supplemented with 3% (w/v) dry milk and incubated with 1 µg/mL 558-6BM IgA1 in blocking buffer. Bound mAb was detected with rabbit anti-human IgA (Dako) followed by HRP-conjugated goat anti-rabbit Ig (SouthernBiotech). Signals were obtained by incubation in West-Pico PLUS substrate (Thermo Fisher Scientific) followed by detection of chemiluminescence on a G:BOX gel doc system (Syngene).

### ELISPOT

To assess secretion of total and antigen-specific antibodies from plasma cells, MultiScreen 96-well filter plates (Sigma) were treated with 35% (v/v) EtOH and coated overnight at 4°C with either 5 µg/mL goat anti-human Ig (SouthernBiotech), 5 µg/mL recombinant human TG2, or 100 µL fixed bacteria in PBS. The membranes were washed twice with PBS and blocked for 2 h with 1% BSA/PBS before addition of duodenal biopsy single-cell suspension or sorted BM plasma cells by applying serial dilutions in RPMI-1640 supplemented with 10% FBS, 10 mM HEPES, pH 7.4, non-essential amino acids, 1 mM sodium pyruvate, and penicillin-streptomycin. After overnight incubation at 37°C, 5% CO_2_, the supernatant was discarded, and the plates were washed extensively with PBS and PBST. Bound antibody was detected by incubation with AP-conjugated goat anti-human IgA (Sigma), goat anti-human IgG (SouthernBiotech) or goat anti-human IgM (SouthernBiotech) in 1% BSA/PBS followed by washing and addition of BCIP/NBT substrate (Bio-Rad). Spots were counted using an ImmunoSpot analyzer (Cellular Technology Limited).

### Detection of Ig modifications by secreted bacterial enzymes

Bacterial strains were grown in 5 mL of BHIYE medium for 24-48 h at 37°C, 5% CO_2_, before the cells were pelleted, and the supernatants were passed through a 0.2 µm filter. 10 µL of supernatant was incubated with 2 µg of purified serum IgA or IgG overnight at 37°C. The samples were then subjected to reducing SDS-PAGE followed by detection of antibody bands with PageBlue protein staining solution (Thermo Fisher Scientific).

## Notes

### Competing Interest Statement

The authors have declared no competing interest.

## REFERENCES

1. Macpherson, A.J., McCoy, K.D., Johansen, F.E., and Brandtzaeg, P. (2008). The immune geography of IgA induction and function. Mucosal Immunol 1, 11–22, 2008. 10.1038/mi.2007.6.

2. Corthesy, B. (2013). Multi-faceted functions of secretory IgA at mucosal surfaces. Front Immunol 4, 185. 10.3389/fimmu.2013.00185.

3. Nakajima, A., Vogelzang, A., Maruya, M., Miyajima, M., Murata, M., Son, A., Kuwahara, T., Tsuruyama, T., Yamada, S., Matsuura, M., et al. (2018). IgA regulates the composition and metabolic function of gut microbiota by promoting symbiosis between bacteria. J Exp Med 215, 2019–2034. 10.1084/jem.20180427.

4. Joglekar, P., Ding, H., Canales-Herrerias, P., Pasricha, P.J., Sonnenburg, J.L., and Peterson, D.A. (2019). Intestinal IgA regulates expression of a fructan polysaccharide utilization locus in colonizing gut commensal bacteroides thetaiotaomicron. mBio 10. 10.1128/mBio.02324-19.

5. Donaldson, G.P., Ladinsky, M.S., Yu, K.B., Sanders, J.G., Yoo, B.B., Chou, W.C., Conner, M.E., Earl, A.M., Knight, R., Bjorkman, P.J., and Mazmanian, S.K. (2018). Gut microbiota utilize immunoglobulin A for mucosal colonization. Science 360, 795–800. 10.1126/science.aaq0926.

6. Benner, R., Hijmans, W., and Haaijman, J.J. (1981). The bone marrow: the major source of serum immunoglobulins, but still a neglected site of antibody formation. Clin Exp Immunol 46, 1–8.

7. Lemke, A., Kraft, M., Roth, K., Riedel, R., Lammerding, D., and Hauser, A.E. (2016). Long-lived plasma cells are generated in mucosal immune responses and contribute to the bone marrow plasma cell pool in mice. Mucosal Immunol 9, 83–97. 10.1038/mi.2015.38.

8. Bemark, M., Hazanov, H., Stromberg, A., Komban, R., Holmqvist, J., Koster, S., Mattsson, J., Sikora, P., Mehr, R., and Lycke, N.Y. (2016). Limited clonal relatedness between gut IgA plasma cells and memory B cells after oral immunization. Nat Commun 7, 12698. 10.1038/ncomms12698.

9. Wilmore, J.R., Gaudette, B.T., Gomez Atria, D., Rosenthal, R.L., Reiser, S.K., Meng, W., Rosenfeld, A.M., Luning Prak, E.T., and Allman, D. (2021). IgA plasma cells are long-lived residents of gut and bone marrow that express isotype- and tissue-specific gene expression patterns. Front Immunol 12, 791095. 10.3389/fimmu.2021.791095.

10. Palm, N.W., de Zoete, M.R., Cullen, T.W., Barry, N.A., Stefanowski, J., Hao, L., Degnan, P.H., Hu, J., Peter, I., Zhang, W., et al. (2014). Immunoglobulin A coating identifies colitogenic bacteria in inflammatory bowel disease. Cell 158, 1000–1010. 10.1016/j.cell.2014.08.006.

11. Bunker, J.J., Flynn, T.M., Koval, J.C., Shaw, D.G., Meisel, M., McDonald, B.D., Ishizuka, I.E., Dent, A.L., Wilson, P.C., Jabri, B., et al. (2015). Innate and adaptive humoral responses coat distinct commensal bacteria with immunoglobulin A. Immunity 43, 541–553. 10.1016/j.immuni.2015.08.007.

12. Sterlin, D., Fadlallah, J., Adams, O., Fieschi, C., Parizot, C., Dorgham, K., Rajkumar, A., Autaa, G., El-Kafsi, H., Charuel, J.L., et al. (2020). Human IgA binds a diverse array of commensal bacteria. J Exp Med 217. 10.1084/jem.20181635.

13. Macpherson, A.J., Gatto, D., Sainsbury, E., Harriman, G.R., Hengartner, H., and Zinkernagel, R.M. (2000). A primitive T cell-independent mechanism of intestinal mucosal IgA responses to commensal bacteria. Science 288, 2222–2226.

14. Macpherson, A.J., Geuking, M.B., Slack, E., Hapfelmeier, S., and McCoy, K.D. (2012). The habitat, double life, citizenship, and forgetfulness of IgA. Immunol Rev 245, 132–146. 10.1111/j.1600-065X.2011.01072.x.

15. Wilmore, J.R., Gaudette, B.T., Gomez Atria, D., Hashemi, T., Jones, D.D., Gardner, C.A., Cole, S.D., Misic, A.M., Beiting, D.P., and Allman, D. (2018). Commensal microbes induce serum IgA responses that protect against polymicrobial sepsis. Cell Host Microbe 23, 302–311 e303. 10.1016/j.chom.2018.01.005.

16. Fadlallah, J., Sterlin, D., Fieschi, C., Parizot, C., Dorgham, K., El Kafsi, H., Autaa, G., Ghillani-Dalbin, P., Juste, C., Lepage, P., et al. (2019). Synergistic convergence of microbiota-specific systemic IgG and secretory IgA. J Allergy Clin Immunol 143, 1575–1585 e1574. 10.1016/j.jaci.2018.09.036.

17. Yang, C., Chen-Liaw, A., Spindler, M.P., Tortorella, D., Moran, T.M., Cerutti, A., and Faith, J.J. (2022). Immunoglobulin A antibody composition is sculpted to bind the self gut microbiome. Sci Immunol 7, eabg3208. 10.1126/sciimmunol.abg3208.

18. Kett, K., Brandtzaeg, P., Radl, J., and Haaijman, J.J. (1986). Different subclass distribution of IgA-producing cells in human lymphoid organs and various secretory tissues. J Immunol 136, 3631–3635.

19. Brandtzaeg, P., and Johansen, F.E. (2005). Mucosal B cells: phenotypic characteristics, transcriptional regulation, and homing properties. Immunol Rev 206, 32–63. 10.1111/j.0105-2896.2005.00283.x.

20. Arroyo Vazquez, J.A., Henning, C., Park, P.O., and Bergstrom, M. (2020). Bacterial colonization of the stomach and duodenum in a Swedish population with and without proton pump inhibitor treatment. JGH Open 4, 405–409. 10.1002/jgh3.12265.

21. Caminero, A., Galipeau, H.J., McCarville, J.L., Johnston, C.W., Bernier, S.P., Russell, A.K., Jury, J., Herran, A.R., Casqueiro, J., Tye-Din, J.A., et al. (2016). Duodenal bacteria from patients with celiac disease and healthy subjects distinctly affect gluten breakdown and immunogenicity. Gastroenterology 151, 670–683. 10.1053/j.gastro.2016.06.041.

22. Collado, M.C., Donat, E., Ribes-Koninckx, C., Calabuig, M., and Sanz, Y. (2009). Specific duodenal and faecal bacterial groups associated with paediatric coeliac disease. J Clin Pathol 62, 264–269. 10.1136/jcp.2008.061366.

23. D’Argenio, V., Casaburi, G., Precone, V., Pagliuca, C., Colicchio, R., Sarnataro, D., Discepolo, V., Kim, S.M., Russo, I., Del Vecchio Blanco, G., et al. (2016). Metagenomics reveals dysbiosis and a potentially pathogenic N. flavescens strain in duodenum of adult celiac patients. Am J Gastroenterol 111, 879–890. 10.1038/ajg.2016.95.

24. Herran, A.R., Perez-Andres, J., Caminero, A., Nistal, E., Vivas, S., Ruiz de Morales, J.M., and Casqueiro, J. (2017). Gluten-degrading bacteria are present in the human small intestine of healthy volunteers and celiac patients. Res Microbiol 168, 673–684. 10.1016/j.resmic.2017.04.008.

25. Nelson, D.P., and Mata, L.J. (1970). Bacterial flora associated with the human gastrointestinal mucosa. Gastroenterology 58, 56–61.

26. Nistal, E., Caminero, A., Herran, A.R., Perez-Andres, J., Vivas, S., Ruiz de Morales, J.M., Saenz de Miera, L.E., and Casqueiro, J. (2016). Study of duodenal bacterial communities by 16S rRNA gene analysis in adults with active celiac disease vs non-celiac disease controls. J Appl Microbiol 120, 1691–1700. 10.1111/jam.13111.

27. Sanchez, E., Donat, E., Ribes-Koninckx, C., Fernandez-Murga, M.L., and Sanz, Y. (2013). Duodenal-mucosal bacteria associated with celiac disease in children. Appl Environ Microbiol 79, 5472–5479. 10.1128/AEM.00869-13.

28. Lycke, N.Y., and Bemark, M. (2017). The regulation of gut mucosal IgA B-cell responses: recent developments. Mucosal Immunol 10, 1361–1374. 10.1038/mi.2017.62.

29. Pabst, O., and Slack, E. (2020). IgA and the intestinal microbiota: the importance of being specific. Mucosal Immunol 13, 12–21. 10.1038/s41385-019-0227-4.

30. Iversen, R., and Sollid, L.M. (2023). The immunobiology and pathogenesis of celiac disease. Annu Rev Pathol 18, 47–70. 10.1146/annurev-pathmechdis-031521-032634.

31. Iversen, R., Snir, O., Stensland, M., Kroll, J.E., Steinsbo, O., Korponay-Szabo, I.R., Lundin, K.E.A., de Souza, G.A., and Sollid, L.M. (2017). Strong clonal relatedness between serum and gut IgA despite different plasma cell origins. Cell Rep 20, 2357–2367. 10.1016/j.celrep.2017.08.036.

32. Stoeckius, M., Hafemeister, C., Stephenson, W., Houck-Loomis, B., Chattopadhyay, P.K., Swerdlow, H., Satija, R., and Smibert, P. (2017). Simultaneous epitope and transcriptome measurement in single cells. Nat Methods 14, 865–868. 10.1038/nmeth.4380.

33. Lindeman, I., Hoydahl, L.S., Christophersen, A., Risnes, L.F., Jahnsen, J., Lundin, K.E.A., Sollid, L.M., and Iversen, R. (2024). Generation of circulating autoreactive pre-plasma cells fueled by naive B cells in celiac disease. Cell Rep 43, 114045. 10.1016/j.celrep.2024.114045.

34. Vidarsson, G., Dekkers, G., and Rispens, T. (2014). IgG subclasses and allotypes: from structure to effector functions. Front Immunol 5, 520. 10.3389/fimmu.2014.00520.

35. Delacroix, D.L., Dive, C., Rambaud, J.C., and Vaerman, J.P. (1982). IgA subclasses in various secretions and in serum. Immunology 47, 383–385.

36. Su, W., Gordon, J.N., Barone, F., Boursier, L., Turnbull, W., Mendis, S., Dunn-Walters, D.K., and Spencer, J. (2008). Lambda light chain revision in the human intestinal IgA response. J Immunol 181, 1264–1271. 10.4049/jimmunol.181.2.1264.

37. Nutt, S.L., Hodgkin, P.D., Tarlinton, D.M., and Corcoran, L.M. (2015). The generation of antibody-secreting plasma cells. Nat Rev Immunol 15, 160–171. 10.1038/nri3795.

38. Winkelmann, R., Sandrock, L., Porstner, M., Roth, E., Mathews, M., Hobeika, E., Reth, M., Kahn, M.L., Schuh, W., and Jack, H.M. (2011). B cell homeostasis and plasma cell homing controlled by Kruppel-like factor 2. Proc Natl Acad Sci U S A 108, 710–715. 10.1073/pnas.1012858108.

39. Kunkel, E.J., Kim, C.H., Lazarus, N.H., Vierra, M.A., Soler, D., Bowman, E.P., and Butcher, E.C. (2003). CCR10 expression is a common feature of circulating and mucosal epithelial tissue IgA Ab-secreting cells. J Clin Invest 111, 1001–1010. 10.1172/JCI17244.

40. Moratz, C., Hayman, J.R., Gu, H., and Kehrl, J.H. (2004). Abnormal B-cell responses to chemokines, disturbed plasma cell localization, and distorted immune tissue architecture in Rgs1-/- mice. Mol Cell Biol 24, 5767–5775. 10.1128/MCB.24.13.5767-5775.2004.

41. Gibbons, D.L., Abeler-Dorner, L., Raine, T., Hwang, I.Y., Jandke, A., Wencker, M., Deban, L., Rudd, C.E., Irving, P.M., Kehrl, J.H., and Hayday, A.C. (2011). Cutting Edge: Regulator of G protein signaling-1 selectively regulates gut T cell trafficking and colitic potential. J Immunol 187, 2067–2071. 10.4049/jimmunol.1100833.

42. Yoon, S.O., Zhang, X., Lee, I.Y., Spencer, N., Vo, P., and Choi, Y.S. (2013). CD9 is a novel marker for plasma cell precursors in human germinal centers. Biochem Biophys Res Commun 431, 41–46. 10.1016/j.bbrc.2012.12.102.

43. Shao, Y., Kim, S.Y., Shin, D., Kim, M.S., Suh, H.W., Piao, Z.H., Jeong, M., Lee, S.H., Yoon, S.R., Lim, B.H., et al. (2010). TXNIP regulates germinal center generation by suppressing BCL-6 expression. Immunol Lett 129, 78–84. 10.1016/j.imlet.2010.02.002.

44. Ziegler, S.F., Ramsdell, F., and Alderson, M.R. (1994). The activation antigen CD69. Stem Cells 12, 456–465. 10.1002/stem.5530120502.

45. Chiles, T.C., and Rothstein, T.L. (1992). Surface Ig receptor-induced nuclear AP-1-dependent gene expression in B lymphocytes. J Immunol 149, 825–831.

46. Pinto, D., Montani, E., Bolli, M., Garavaglia, G., Sallusto, F., Lanzavecchia, A., and Jarrossay, D. (2013). A functional BCR in human IgA and IgM plasma cells. Blood 121, 4110–4114. 10.1182/blood-2012-09-459289.

47. Xu, A.Q., Barbosa, R.R., and Calado, D.P. (2020). Genetic timestamping of plasma cells in vivo reveals tissue-specific homeostatic population turnover. Elife 9. 10.7554/eLife.59850.

48. Asmar, A.T., and Collet, J.F. (2018). Lpp, the Braun lipoprotein, turns 50-major achievements and remaining issues. FEMS Microbiol Lett 365. 10.1093/femsle/fny199.

49. Kornfeld, S.J., and Plaut, A.G. (1981). Secretory immunity and the bacterial IgA proteases. Rev Infect Dis 3, 521–534. 10.1093/clinids/3.3.521.

50. Mulks, M.H., Kornfeld, S.J., and Plaut, A.G. (1980). Specific proteolysis of human IgA by Streptococcus pneumoniae and Haemophilus influenzae. J Infect Dis 141, 450–456. 10.1093/infdis/141.4.450.

51. Poulsen, K., Reinholdt, J., Jespersgaard, C., Boye, K., Brown, T.A., Hauge, M., and Kilian, M. (1998). A comprehensive genetic study of streptococcal immunoglobulin A1 proteases: evidence for recombination within and between species. Infect Immun 66, 181–190. 10.1128/IAI.66.1.181-190.1998.

52. Bek-Thomsen, M., Poulsen, K., and Kilian, M. (2012). Occurrence and evolution of the paralogous zinc metalloproteases IgA1 protease, ZmpB, ZmpC, and ZmpD in Streptococcus pneumoniae and related commensal species. mBio 3. 10.1128/mBio.00303-12.

53. Plaut, A.G., Wistar, R., Jr., and Capra, J.D. (1974). Differential susceptibility of human IgA immunoglobulins to streptococcal IgA protease. J Clin Invest 54, 1295–1300. 10.1172/JCI107875.

54. Reinholdt, J., Tomana, M., Mortensen, S.B., and Kilian, M. (1990). Molecular aspects of immunoglobulin A1 degradation by oral streptococci. Infect Immun 58, 1186–1194. 10.1128/iai.58.5.1186-1194.1990.

55. Kilian, M., and Tettelin, H. (2019). Identification of virulence-associated properties by comparative genome analysis of Streptococcus pneumoniae, S. pseudopneumoniae, S. mitis, three S. oralis subspecies, and S. infantis. mBio 10. 10.1128/mBio.01985-19.

56. Lomholt, J.A., and Kilian, M. (2000). Immunoglobulin A1 protease activity in Gemella haemolysans. J Clin Microbiol 38, 2760–2762. 10.1128/JCM.38.7.2760-2762.2000.

57. Frandsen, E.V. (1994). Carbohydrate depletion of immunoglobulin A1 by oral species of gram-positive rods. Oral Microbiol Immunol 9, 352–358. 10.1111/j.1399-302x.1994.tb00285.x.

58. Benckert, J., Schmolka, N., Kreschel, C., Zoller, M.J., Sturm, A., Wiedenmann, B., and Wardemann, H. (2011). The majority of intestinal IgA+ and IgG+ plasmablasts in the human gut are antigen-specific. J Clin Invest 121, 1946–1955. 10.1172/JCI44447.

59. Kabbert, J., Benckert, J., Rollenske, T., Hitch, T.C.A., Clavel, T., Cerovic, V., Wardemann, H., and Pabst, O. (2020). High microbiota reactivity of adult human intestinal IgA requires somatic mutations. J Exp Med 217. 10.1084/jem.20200275.

60. Dunn-Walters, D.K., Boursier, L., and Spencer, J. (1997). Hypermutation, diversity and dissemination of human intestinal lamina propria plasma cells. Eur J Immunol 27, 2959–2964. 10.1002/eji.1830271131.

61. Rollenske, T., Szijarto, V., Lukasiewicz, J., Guachalla, L.M., Stojkovic, K., Hartl, K., Stulik, L., Kocher, S., Lasitschka, F., Al-Saeedi, M., et al. (2018). Cross-specificity of protective human antibodies against Klebsiella pneumoniae LPS O-antigen. Nat Immunol 19, 617–624. 10.1038/s41590-018-0106-2.

62. Sollid, L.M., and Iversen, R. (2023). Tango of B cells with T cells in the making of secretory antibodies to gut bacteria. Nat Rev Gastroenterol Hepatol 20, 120–128. 10.1038/s41575-022-00674-y.

63. Brandtzaeg, P., Farstad, I.N., and Haraldsen, G. (1999). Regional specialization in the mucosal immune system: primed cells do not always home along the same track. Immunol Today 20, 267–277. 10.1016/s0167-5699(99)01468-1.

64. Lin, M., Du, L., Brandtzaeg, P., and Pan-Hammarstrom, Q. (2014). IgA subclass switch recombination in human mucosal and systemic immune compartments. Mucosal Immunol 7, 511–520. 10.1038/mi.2013.68.

65. Horns, F., Vollmers, C., Croote, D., Mackey, S.F., Swan, G.E., Dekker, C.L., Davis, M.M., and Quake, S.R. (2016). Lineage tracing of human B cells reveals the in vivo landscape of human antibody class switching. Elife 5, e16578. 10.7554/eLife.16578.

66. Fenton, T.M., Jorgensen, P.B., Niss, K., Rubin, S.J.S., Morbe, U.M., Riis, L.B., Da Silva, C., Plumb, A., Vandamme, J., Jakobsen, H.L., et al. (2020). Immune profiling of human gut-associated lymphoid tissue identifies a role for isolated lymphoid follicles in priming of region-specific immunity. Immunity 52, 557–570 e556. 10.1016/j.immuni.2020.02.001.

67. Tejedor Vaquero, S., Neuman, H., Comerma, L., Marcos-Fa, X., Corral-Vazquez, C., Uzzan, M., Pybus, M., Segura-Garzon, D., Guerra, J., Perruzza, L., et al. (2024). Immunomolecular and reactivity landscapes of gut IgA subclasses in homeostasis and inflammatory bowel disease. J Exp Med 221. 10.1084/jem.20230079.

68. Lindner, C., Wahl, B., Fohse, L., Suerbaum, S., Macpherson, A.J., Prinz, I., and Pabst, O. (2012). Age, microbiota, and T cells shape diverse individual IgA repertoires in the intestine. J Exp Med 209, 365–377. 10.1084/jem.20111980.

69. Ramirez, S.I., Faraji, F., Hills, L.B., Lopez, P.G., Goodwin, B., Stacey, H.D., Sutton, H.J., Hastie, K.M., Saphire, E.O., Kim, H.J., et al. (2024). Immunological memory diversity in the human upper airway. Nature 632, 630–636. 10.1038/s41586-024-07748-8.

70. Liu, J., Stoler-Barak, L., Hezroni-Bravyi, H., Biram, A., Lebon, S., Davidzohn, N., Kedmi, M., Chemla, M., Pilzer, D., Cohen, M., et al. (2024). Turbinate-homing IgA-secreting cells originate in the nasal lymphoid tissues. Nature 632, 637–646. 10.1038/s41586-024-07729-x.

71. Johansen, F.E., Baekkevold, E.S., Carlsen, H.S., Farstad, I.N., Soler, D., and Brandtzaeg, P. (2005). Regional induction of adhesion molecules and chemokine receptors explains disparate homing of human B cells to systemic and mucosal effector sites: dispersion from tonsils. Blood 106, 593–600. 10.1182/blood-2004-12-4630.

72. Quiding-Jarbrink, M., Granstrom, G., Nordstrom, I., Holmgren, J., and Czerkinsky, C. (1995). Induction of compartmentalized B-cell responses in human tonsils. Infect Immun 63, 853–857. 10.1128/iai.63.3.853-857.1995.

73. Rudin, A., Johansson, E.L., Bergquist, C., and Holmgren, J. (1998). Differential kinetics and distribution of antibodies in serum and nasal and vaginal secretions after nasal and oral vaccination of humans. Infect Immun 66, 3390–3396. 10.1128/IAI.66.7.3390-3396.1998.

74. Currie, E.G., Coburn, B., Porfilio, E.A., Lam, P., Rojas, O.L., Novak, J., Yang, S., Chowdhury, R.B., Ward, L.A., Wang, P.W., et al. (2022). Immunoglobulin A nephropathy is characterized by anticommensal humoral immune responses. JCI Insight 7. 10.1172/jci.insight.141289.

75. Nguyen, M.T., Matsuo, M., Niemann, S., Herrmann, M., and Gotz, F. (2020). Lipoproteins in gram-positive bacteria: abundance, function, fitness. Front Microbiol 11, 582582. 10.3389/fmicb.2020.582582.

76. Jung, J., Choe, J., Li, L., and Choi, Y.S. (2000). Regulation of CD27 expression in the course of germinal center B cell differentiation: the pivotal role of IL-10. Eur J Immunol 30, 2437–2443. 10.1002/1521-4141(2000)30:8<2437::AID-IMMU2437>3.0.CO;2-M.

77. Victora, G.D., and Nussenzweig, M.C. (2022). Germinal centers. Annu Rev Immunol 40, 413–442. 10.1146/annurev-immunol-120419-022408.

78. Robinson, M.J., Webster, R.H., and Tarlinton, D.M. (2020). How intrinsic and extrinsic regulators of plasma cell survival might intersect for durable humoral immunity. Immunol Rev 296, 87–103. 10.1111/imr.12895.

79. Sulkanen, S., Halttunen, T., Laurila, K., Kolho, K.L., Korponay-Szabo, I.R., Sarnesto, A., Savilahti, E., Collin, P., and Maki, M. (1998). Tissue transglutaminase autoantibody enzyme-linked immunosorbent assay in detecting celiac disease. Gastroenterology 115, 1322–1328, 1998. 10.1016/s0016-5085(98)70008-3.

80. Royle, L., Roos, A., Harvey, D.J., Wormald, M.R., van Gijlswijk-Janssen, D., Redwan el, R.M., Wilson, I.A., Daha, M.R., Dwek, R.A., and Rudd, P.M. (2003). Secretory IgA N- and O-glycans provide a link between the innate and adaptive immune systems. J Biol Chem 278, 20140–20153. 10.1074/jbc.M301436200.

81. Li, H., Limenitakis, J.P., Greiff, V., Yilmaz, B., Scharen, O., Urbaniak, C., Zund, M., Lawson, M.A.E., Young, I.D., Rupp, S., et al. (2020). Mucosal or systemic microbiota exposures shape the B cell repertoire. Nature 584, 274–278. 10.1038/s41586-020-2564-6.

82. Rollenske, T., Burkhalter, S., Muerner, L., von Gunten, S., Lukasiewicz, J., Wardemann, H., and Macpherson, A.J. (2021). Parallelism of intestinal secretory IgA shapes functional microbial fitness. Nature 598, 657–661. 10.1038/s41586-021-03973-7.

83. Al-Toma, A., Volta, U., Auricchio, R., Castillejo, G., Sanders, D.S., Cellier, C., Mulder, C.J., and Lundin, K.E.A. (2019). European Society for the Study of Coeliac Disease (ESsCD) guideline for coeliac disease and other gluten-related disorders. United European Gastroenterol J 7, 583–613. 10.1177/2050640619844125.

84. Hao, Y., Stuart, T., Kowalski, M.H., Choudhary, S., Hoffman, P., Hartman, A., Srivastava, A., Molla, G., Madad, S., Fernandez-Granda, C., and Satija, R. (2024). Dictionary learning for integrative, multimodal and scalable single-cell analysis. Nat Biotechnol 42, 293–304. 10.1038/s41587-023-01767-y.

85. Alamyar, E., Giudicelli, V., Li, S., Duroux, P., and Lefranc, M.P. (2012). IMGT/HighV-QUEST: the IMGT® web portal for immunoglobulin (IG) or antibody and T cell receptor (TR) analysis from NGS high throughput and deep sequencing. Immunome Res 8, 26.

86. Lindeman, I., Emerton, G., Mamanova, L., Snir, O., Polanski, K., Qiao, S.W., Sollid, L.M., Teichmann, S.A., and Stubbington, M.J.T. (2018). BraCeR: B-cell-receptor reconstruction and clonality inference from single-cell RNA-seq. Nat Methods 15, 563–565. 10.1038/s41592-018-0082-3.

87. Gupta, N.T., Vander Heiden, J.A., Uduman, M., Gadala-Maria, D., Yaari, G., and Kleinstein, S.H. (2015). Change-O: a toolkit for analyzing large-scale B cell immunoglobulin repertoire sequencing data. Bioinformatics 31, 3356–3358. 10.1093/bioinformatics/btv359.

88. Zhou, C., Osterbye, T., Bach, E., Dahal-Koirala, S., Hoydahl, L.S., Steinsbo, O., Jahnsen, J., Lundin, K.E.A., Buus, S., Sollid, L.M., and Iversen, R. (2022). Focused B cell response to recurring gluten motif with implications for epitope spreading in celiac disease. Cell Rep 41, 111541. 10.1016/j.celrep.2022.111541.

89. Tiller, T., Meffre, E., Yurasov, S., Tsuiji, M., Nussenzweig, M.C., and Wardemann, H. (2008). Efficient generation of monoclonal antibodies from single human B cells by single cell RT-PCR and expression vector cloning. J Immunol Methods 329, 112–124. 10.1016/j.jim.2007.09.017.

90. Iversen, R., Fleur du Pre, M., Di Niro, R., and Sollid, L.M. (2015). Igs as substrates for transglutaminase 2: implications for autoantibody production in celiac disease. J Immunol 195, 5159–5168. 10.4049/jimmunol.1501363.

91. Das, S., Stamnaes, J., Høydahl, L.S., Skagen, C., Lundin, K.E.A., Jahnsen, J., Sollid, L.M., and Iversen, R. (2024). Selective activation of naïve B cells with unique epitope specificity shapes autoantibody formation in celiac disease. J Autoimmun 146, 103241. 10.1016/j.jaut.2024.103241.

92. Stamnaes, J., Cardoso, I., Iversen, R., and Sollid, L.M. (2016). Transglutaminase 2 strongly binds to an extracellular matrix component other than fibronectin via its second C-terminal beta-barrel domain. FEBS J 283, 3994–4010. 10.1111/febs.13907.

