## Supplemental info for "Distinct systemic and gut IgA responses to bacteria of the human upper gastrointestinal tract"

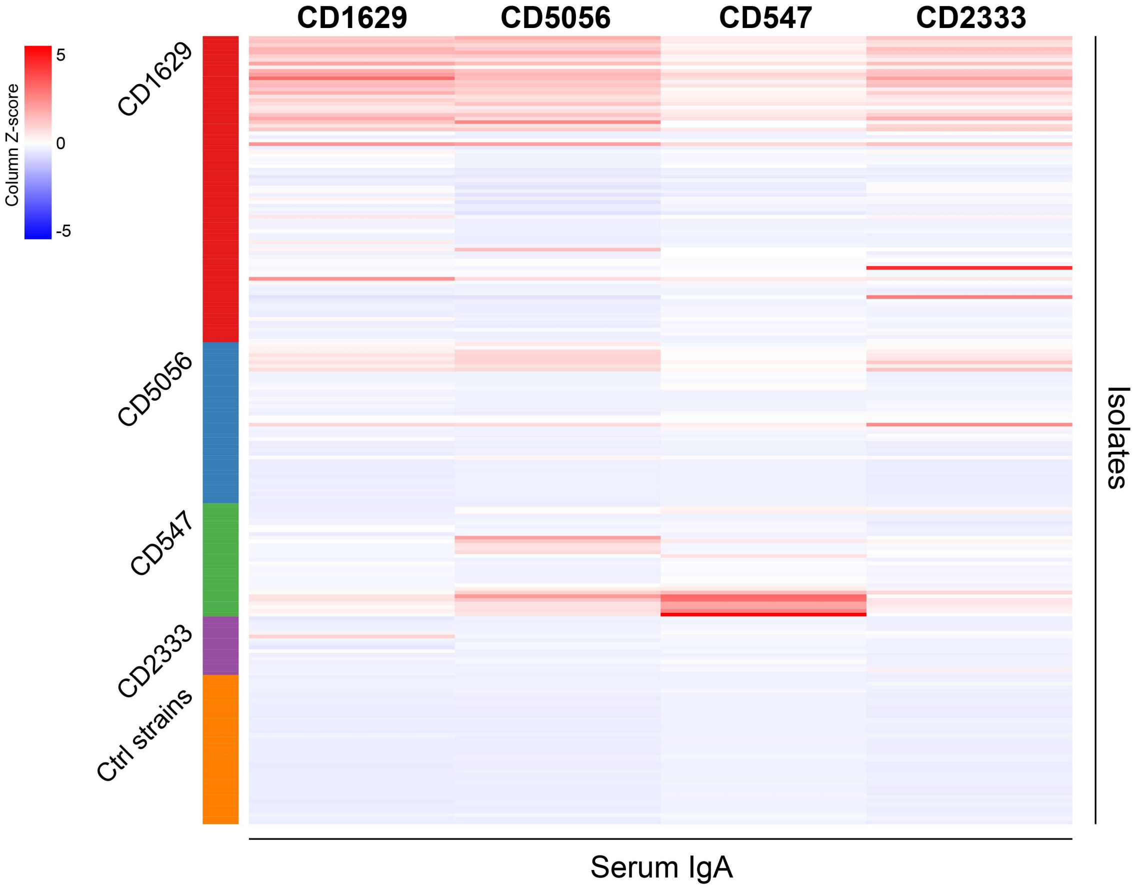

**Figure S1. Reactivity of serum IgA against self- and non-self-bacteria.**
Heat map showing reactivity of serum IgA purified from four donors against individual bacterial strains (n = 216) as determined by flow cytometry. The strains were isolated from duodenal biopsies of the same four donors, or they were non-duodenum reference strains.

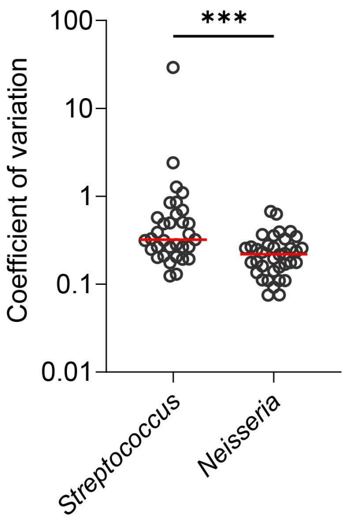

**Figure S2. Variation in antibody responses over time.**
Variation in serum IgA responses of a single donor (CD1629) against individual bacterial strains of the genera *Streptococcus* (n = 33) or *Neisseria* (n = 36). IgA binding was assessed by flow cytometry, and the coefficient of variation was calculated for each strain based on serum samples collected at four discrete time points. Only strains with ∆MFI values >10% of the maximum signal in at least one time point were included. Horizontal lines indicate medians and difference between groups was evaluated by a Mann-Whitney U test. ***$p < 0.001$.

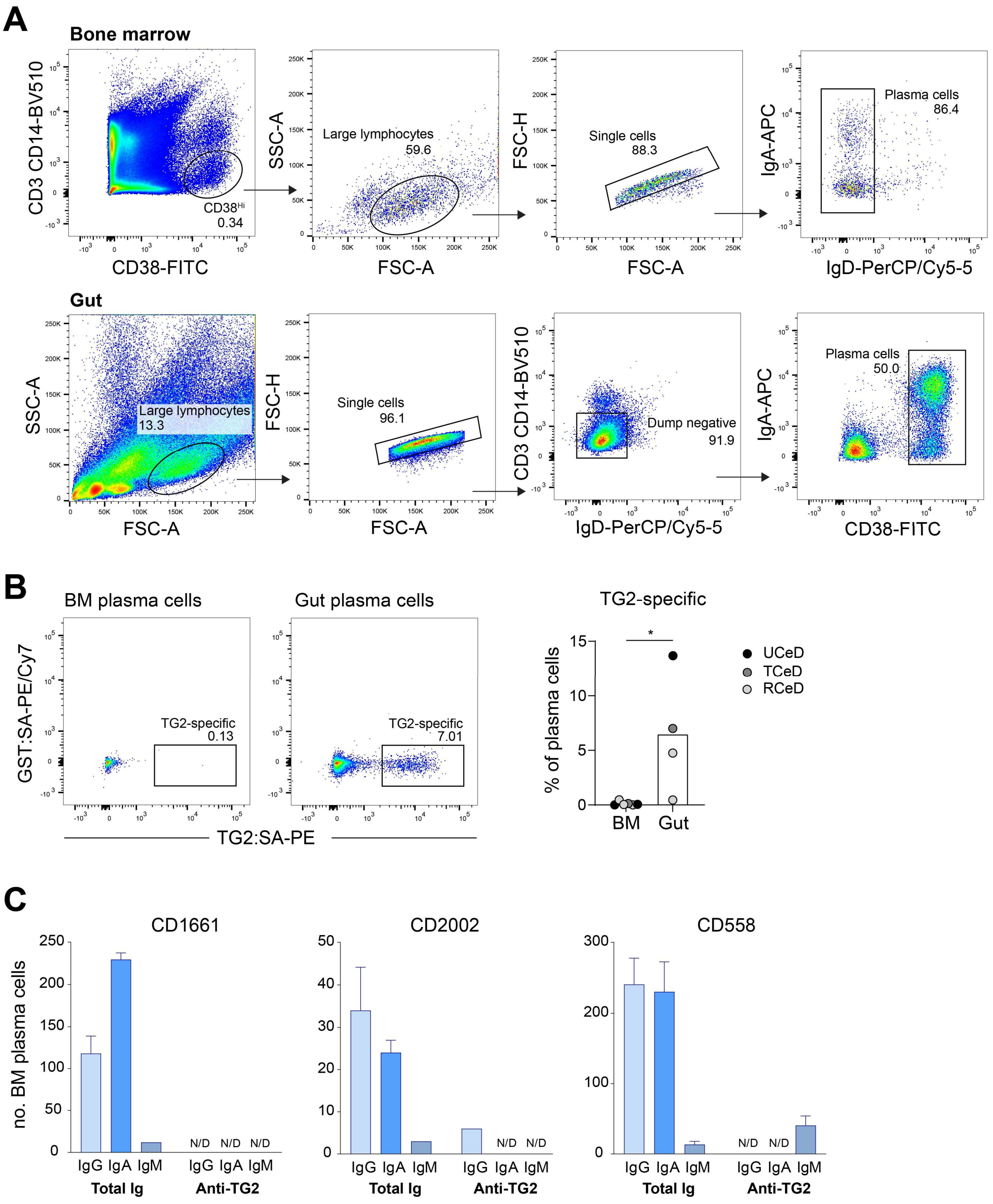

**Figure S3. Detection of TG2-specific plasma cells.**

(A) Gating strategy for identification of plasma cells in bone marrow (BM) aspirates and gut biopsies.
(B) Representative flow cytometry plots and summary data of four donors showing staining of bone marrow (BM) or gut plasma cells with recombinant biotinylated TG2 bound to PE-conjugated streptavidin (SA-PE) in combination with an irrelevant control antigen (glutathione S-transferase; GST). The donors were patients with untreated celiac disease (UCeD), treated celiac disease (TCeD, gluten-free diet) or refractory celiac disease (RCeD). Bar heights indicate means, and difference between groups was evaluated by a paired t test. *$p < 0.05$.
(C) Detection of total Ig-secreting and anti-TG2-secreting BM plasma cells in three celiac disease patients by ELISPOT. Bar heights indicate means, and error bars indicate SEM based on four dilutions of BM aspirates. N/D, not detectable.

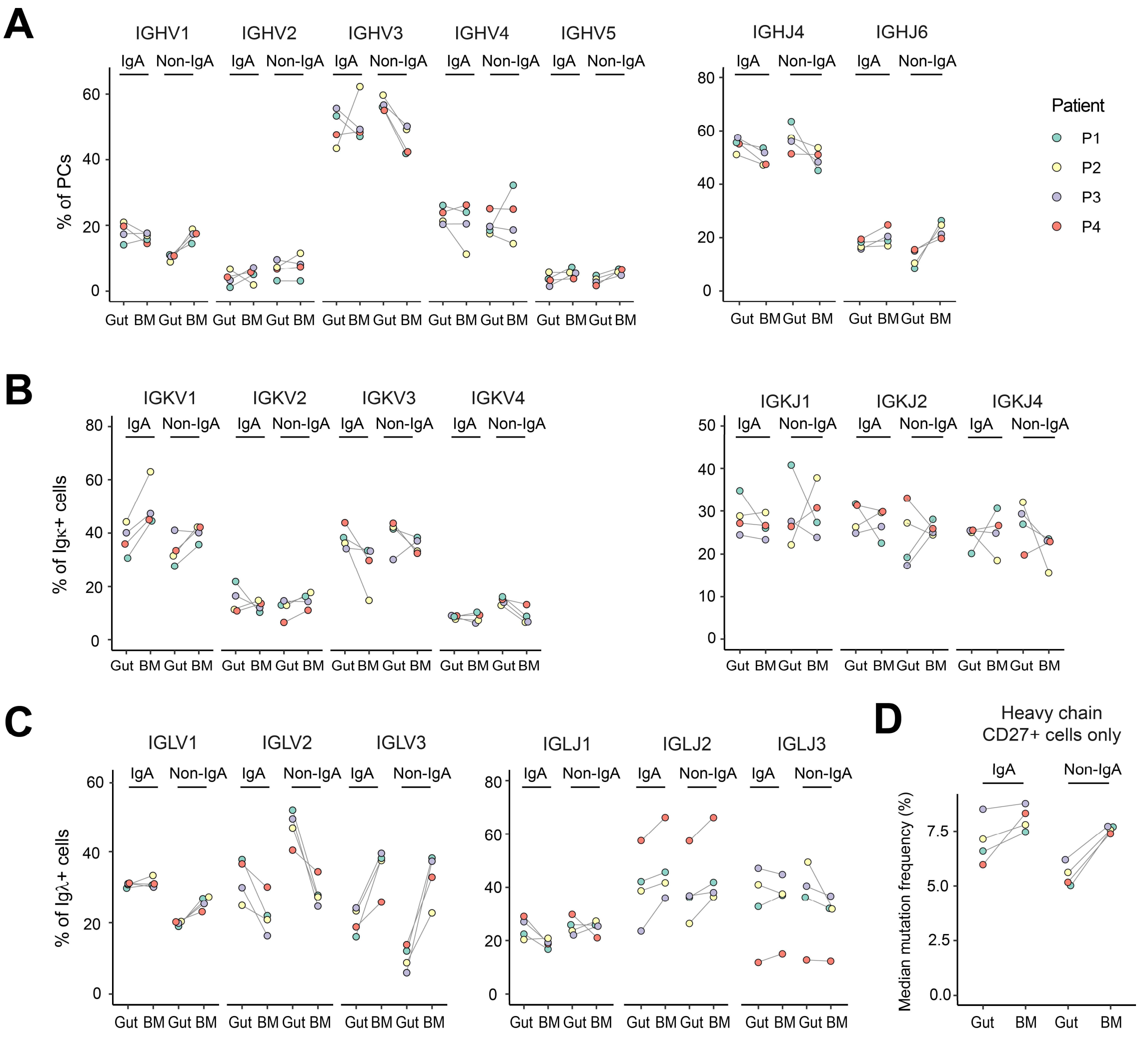

**Figure S4. Comparison of Ig repertoires between gut and BM plasma cells.**
(A-C) Usage of heavy chain (A), kappa light chain (B) and lambda light chain (C) major V- and J-gene families among IgA or non-IgA plasma cells (PCs) isolated from duodenal biopsies or BM aspirates of four patients.
(D) Heavy chain V-gene mutation levels in surface $CD27^{+}$ IgA or non-IgA plasma cells isolated from either gut or BM. In the gut, non-IgA cells are mainly $IgM^{+}$, whereas non-IgA cells in BM are primarily $IgG^{+}$.

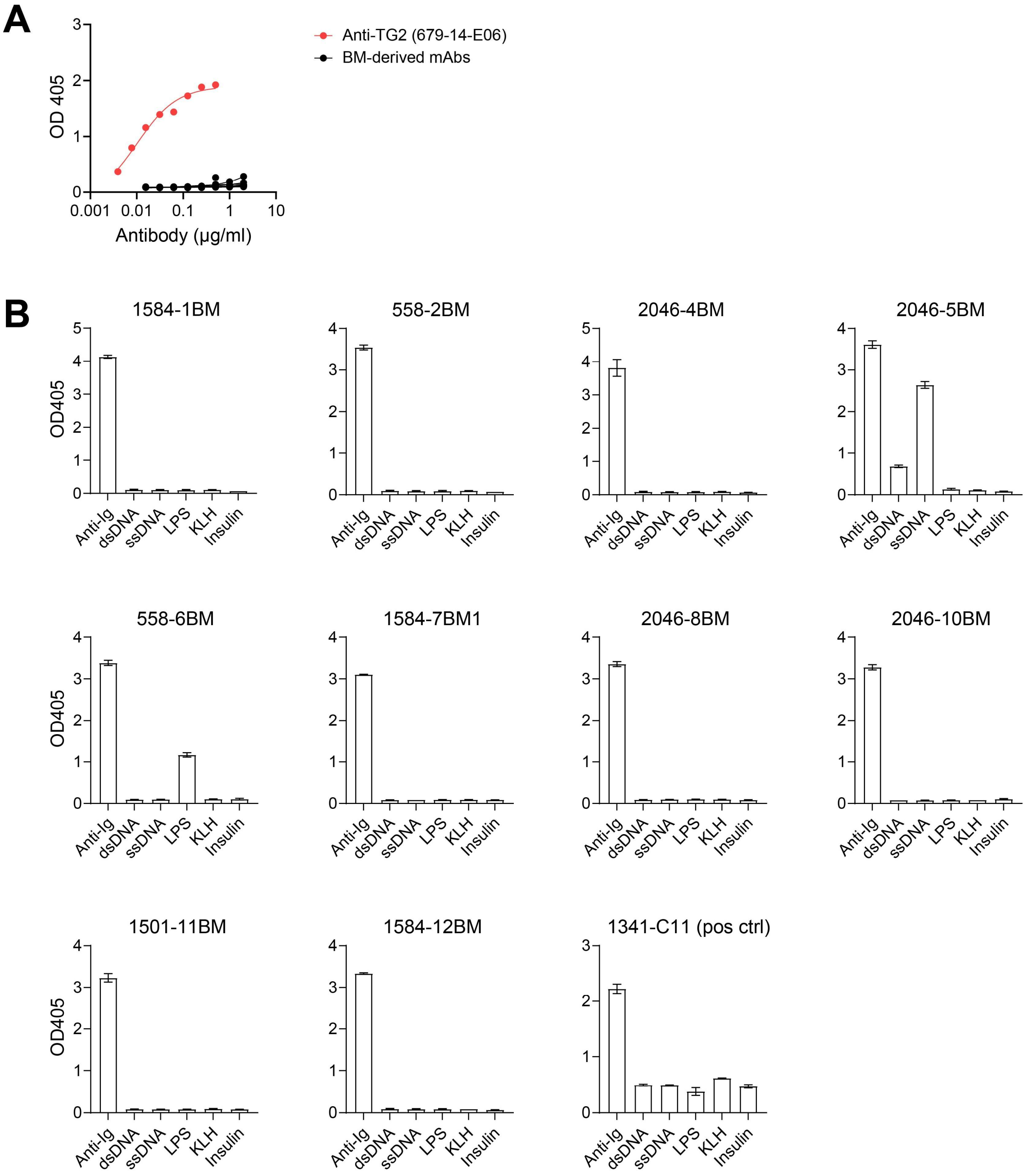

**Figure S5. Reactivity of BM-gut spanning clonotypes**.

(A) ELISA binding curves showing reactivity of mAbs generated from BM plasma cells (n=10) with recombinant human TG2. The previously generated anti-TG2 mAb 679-14-E06 was included as positive control.

(B) Polyreactivity screen for BM-derived mAbs. The following model antigens were tested in ELISA: double-stranded DNA (dsDNA), single-stranded DNA (ssDNA), lipopolysaccharide (LPS), keyhole limpet hemocyanin (KLH), and insulin. A polyclonal goat anti-human Ig antibody was included as positive control antigen. A previously generated polyreactive mAb, 1341-C11, is shown for comparison. All mAbs were expressed in a human IgA1 format (except 1341-C11 which was expressed as IgG1). For polyreactivty screening, mAbs were tested in a concentration of 1 µg/ml. Error bars indicate range of sample duplicates.

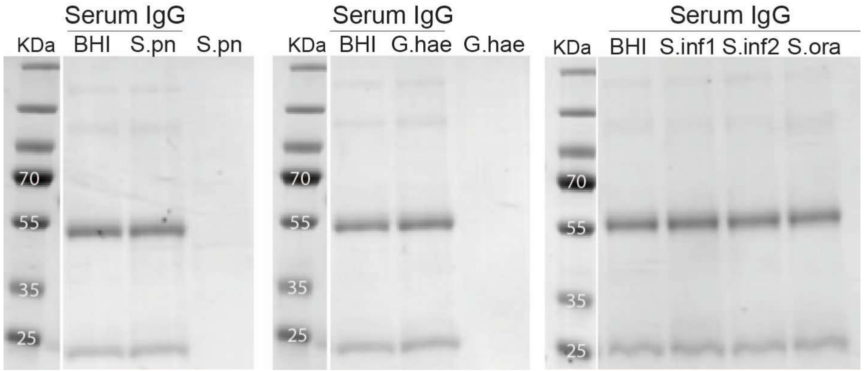

**Figure S6. Effect of bacterial secreted enzymes on IgG.**

Assessment of preserved IgG integrity by SDS-PAGE. Purified serum IgG of a single donor was incubated with culture supernatant of bacteria shown to either degrade IgA: *Streptococcus pneumoniae* (S.pn) and *Gemella haemolysans* (G.hae), or deglycosylate IgA: *Streptococcus infantis* (S.inf1 and S.inf2). Culture supernatants alone and IgG incubated with brain heart infusion (BHI) broth or supernatant of *Streptococcus oralis* (S.ora) were included as controls.

**Table S1. Clinicopathological data of patients donating biological samples**

| | Patient ID | Short name | Gender | Age | Diagnosis[a] |
|---|---|---|---|---|---|
| Isolation of bacteria from gut biopsies | CD1629 | | F | 27 | Chronic non-celiac enteropathy |
| | CD5056 | | F | 60 | UCeD |
| | CD547 | | F | 68 | TCeD |
| | CD2333 | | F | 44 | Not CeD, kidney transplant |
| Collection of BM aspirates | CD558[b] | P1 | M | 71 | TCeD |
| | CD1501[b] | P2 | M | 60 | RCeD |
| | CD1584[b] | P3 | M | 53 | RCeD |
| | CD2046[b] | P4 | F | 43 | UCeD |
| | CD2002 | | F | 42 | UCeD |

[a]UCeD, untreated celiac disease; TCeD, treated celiac disease (gluten-free diet); RCeD, refractory celiac disease (type 2)
[b]Transcriptome of gut and BM plasma cells

**Table S2. Strains isolated from duodenal biopsies of four donors**

| | CD1629 | CD5056 | CD547 | CD2333 |
|---|---|---|---|---|
| *Streptococcus parasanguinis* | 27 | | 7 | 1 |
| *Streptococcus cristatus* | | 8 | | |
| *Streptococcus infantis* | 2 | 5 | | 1 |
| *Streptococcus peroris* | 1 | | | |
| *Streptococcus mitis* | 1 | | | 1 |
| *Streptococcus oralis* | | | | 5 |
| *Streptococcus vestibularis* | | | | 2 |
| *Streptococcus anginosus* | | | 10 | 1 |
| *Streptococcus infantis/mitis* | 1 | | | |
| *Streptococcus pneumoniae/oralis* | 1 | | | |
| *Streptococcus mitis/oralis* | 1 | | | 3 |
| *Streptococcus peroris/oralis* | 1 | | | |
| *Streptococcus vestibularis/salivarius* | | | | 1 |
| Streptococci sp. | | 6 | | |
| Mitis group *Streptococcus* | 4 | | | |
| *Neisseria subflava/perflava* | 2 | | | |
| *Neisseria subflava* | 11 | 2 | | |
| *Neisseria flavescens* | 3 | 2 | | |
| *Neisseria sp.* | 14 | 4 | | |
| *Rothia dentocariosa* | 3 | | | |
| *Rothia mucilaginosa* | 3 | | | |
| *Staphylococcus saprophyticus* | 1 | | | |
| *Staphylococcus warneri* | 1 | | | |
| *Staphylococcus epidermidis* | | 17 | 5 | |
| *Corynebacterium singulare* | 1 | | | |
| *Corynebacterium durum* | 1 | | | |
| *Lactobacillus paracasei* | | | 6 | |
| *Micrococcus luteus* | 2 | | 1 | |
| *Actinomyces oris* | 2 | | | |
| *Gemella sanguinis* | | | | 1 |
| *Gemella haemolysans* | 1 | | | |
| *Candida albicans* | | | 2 | |
| Total | 84 | 44 | 31 | 16 |

**Table S3. List of reference/control strains used in this study**

| Strain | Source/Gift from |
|---|---|
| *Escherichia coli* F-18 | D. Wall, University of Glasgow |
| *Escherichia coli* Nissle 1917 (EcN) | D. Wall, University of Glasgow |
| *Escherichia coli* 217 A1 | D. Walker, University of Glasgow |
| *Escherichia coli* 083 A1 | D. Walker, University of Glasgow |
| *Escherichia coli* LF-82 | D. Walker, University of Glasgow |
| *Escherichia coli* 031 A1 | D. Walker, University of Glasgow |
| *Escherichia coli* 033 A1 | D. Walker, University of Glasgow |
| *Escherichia coli* 035 A1 | D. Walker, University of Glasgow |
| *Enterobacteriaceae* 032 A1 | D. Walker, University of Glasgow |
| *Klebsiella oxytoca* 216 C1 | D. Walker, University of Glasgow |
| *Klebsiella oxytoca* 077 D | D. Walker, University of Glasgow |
| *Enterobacter* sp. 077 C | D. Walker, University of Glasgow |
| *Enterobacter* sp. 216 D1 | D. Walker, University of Glasgow |
| *Klebsiela pneumoniae* TE10 | D. Walker, University of Glasgow |
| *Klebsiela pneumoniae* TE11 | D. Walker, University of Glasgow |
| *Salmonella enterica* subsp. *enterica* serovar Typhimurium SL1344 | D. Wall, University of Glasgow |
| *Salmonella enterica* subsp. *enterica* serovar Typhimurium SL7207 | D. Wall, University of Glasgow |
| *Bacteroides fragilis* NCTC 9343/ATCC 25285 | D. Wall, University of Glasgow |
| *Staphylococcus aureus* Newman | T. Foster, University of Dublin |
| *Staphylococcus aureus* Newman Δ *spa* Δ *sbi* | T. Foster, University of Dublin |
| *Staphylococcus epidermidis* BM96183 | J. Penades, Imperial College London |
| *Staphylococcus epidermidis* BM96184 | J. Penades, Imperial College London |
| *Streptococcus pneumoniae* Pn-20 | J. Penades, Imperial College London |
| *Streptococcus pneumoniae* Hungary 19A-6 (ATCC 700673) | J. Penades, Imperial College London |
| *Streptococcus pyogenes* ATCC 700294 | J. Penades, Imperial College London |
| *Streptococcus pyogenes* ATCC BAA-1633 | J. Penades, Imperial College London |
| *Listeria monocytogenes* SK1351 | J. Penades, Imperial College London |
| *Listeria monocytogenes* EGDE | J. Penades, Imperial College London |
| *Lactococcus lactis* sub. *lactis* IL1403 | J. Penades, Imperial College London |
| *Lactococcus lactis* sub. *cremoris* MG1363 | J. Penades, Imperial College London |
| *Bacillus thuringiensis* HD 395 | J. Penades, Imperial College London |
| *Bacillus thuringiensis* 4Q1 | J. Penades, Imperial College London |
| *Bacillus thuringiensis* ATCC 35646 | J. Penades, Imperial College London |
| *Enterococcus faecalis* NCTC 12201 | D. Walker, University of Glasgow |
| *Enterococcus faecalis* NCTC 12203 | D. Walker, University of Glasgow |
| *Enterococcus faecium* NCTC 12202 | D. Walker, University of Glasgow |
| *Enterococcus faecium* NCTC 13923 | D. Walker, University of Glasgow |
| *Enterococcus faecium* NCTC 12204 | D. Walker, University of Glasgow |
| *Enterococcus casseliflavus* NCTC 12361 | D. Walker, University of Glasgow |
| *Clostridium clostridioforme* NCTC 7155 | D. Wall, University of Glasgow |
| *Clostridiodes difficile* R20291 | G. Douce, University of Glasgow |
| *Bifidobacterium animalis* DSM 26074 | DSMZ, Leibniz Institute |

**Table S4. Sequence properties of mAbs generated from BM-gut spanning clonotypes**

| mAb ID | No. cells[a] | Isotype | V gene | D gene | J gene | Junction | CDR3 length | Mutations[b] |
|---|---|---|---|---|---|---|---|---|
| | BM | BM | IGHV | IGHD | IGHJ | H chain | H chain | H chain |
| | Gut | Gut | IGKV/IGLV | | IGKJ/IGLJ | L chain | L chain | L chain |
| 1584-1BM | 1 | IgA2 | HV3-72*01 | HD3-22*01 | HJ4*02 | CAREGFYDISGVDHW | 13 | 26/6 |
| | 12 | IgA2(7), IgA1(1), IgM(4) | KV2-30*01 or KV2-30*02 | | KJ3*01 | CLQGSHWPFTF | 9 | 14/6 |
| 558-2BM | 1 | IgA2 | HV3-74*01 | HD2-15*01 | HJ5*01 or HJ5*02 | CARDVGGRSSFW | 10 | 9/4 |
| | 7 | IgA2 | KV2-28*01 or KV2D-28*01 | | KJ2*01 | CVQALQTPYTF | 9 | 10/1 |
| 2046-4BM | 1 | IgA1 | HV3-9*04 | HD5-18*01 | HJ6*04 | CTKDLQPGGADVW | 11 | 27/5 |
| | 3 | IgA1 | KV2-30*01 | | KJ2*01 | CMQGTHWPYTF | 9 | 4/0 |
| 2046-5BM | 1 | IgA1 | HV3-7*01 | HD5-12*01 | HJ3*02 | CARAITHGFDMG | 10 | 11/2 |
| | 4 | IgA1(2), IgA2(1), IgA[c](1) | LV2-23*02 | | LJ1*01 | CCSYAGTWYVF | 9 | 15/3 |
| 558-6BM | 1 | IgA2 | HV3-15*01 | HD2-15*01 | HJ4*02 | CTTERDCSGGSCLHNW | 14 | 9/3 |
| | 2 | IgA2 | KV1-5*03 | | KJ2*01 | CQQYYSFHTF | 8 | 3/3 |
| 1584-7BM1 | 3 | IgA1 | HV1-46*01 | HD2-2*02 | HJ6*02 | CARQALGYCSSTSCYKSGMDVW | 20 | 5/0 |
| | 1 | IgA1 | KV3-20*01 | | KJ4*01 | CQQYGSSPLTF | 9 | 5/1 |
| 2046-8BM | 1 | IgG2 | HV3-30*20 | HD5-12*01 | HJ4*02 | CVKENSDYFFDYW | 11 | 19/3 |
| | 2 | IgA2 | KV3-11*01 | | KJ4*01 | CQQRTVWLRTF | 9 | 10/0 |
| 2046-10BM | 1 | IgA2 | HV2-5*02 | HD4-23*01 | HJ4*02 | CAHAYRNNWYIRYW | 12 | 12/1 |
| | 2 | IgA2 | KV4-1*01 | | KJ4*01 | CQQYYDAPLTF | 9 | 5/0 |
| 1501-11BM | 1 | IgA2 | HV3-49*03 | HD6-19*01 | HJ3*02 | CTRGQWLSSWALDIW | 13 | 7/2 |
| | 2 | IgA2 | KV2-24*01 | | KJ2*01 or KJ2*02 | CMQATQFPHSL | 9 | 1/4 |
| 1584-12BM | 1 | IgG1 | HV4-61*01 or HV4-61*03 | HD6-19*01 | HJ6*02 | CAREICRFSGAWDAFGDPGLDVW | 21 | 15/14 |
| | 2 | IgG1 | KV1-17*01 | | KJ5*01 | CLQHNSYPLTF | 9 | 9/2 |

[a]Number of cells in each compartment belonging to the same clonotype

[b]Replacement/silent nt substitutions in V gene

[c]Unknown subclass
